# Computationally efficient framework for diagnosing, understanding, and predicting biphasic population growth

**DOI:** 10.1101/2022.07.27.501797

**Authors:** Ryan J. Murphy, Oliver J. Maclaren, Alivia R. Calabrese, Patrick B. Thomas, David J. Warne, Elizabeth D. Williams, Matthew J. Simpson

## Abstract

Throughout the life sciences, biological populations undergo multiple phases of growth, often referred to as *biphasic growth* for the commonly-encountered situation involving two phases. Biphasic population growth occurs over a massive range of spatial and temporal scales, ranging from microscopic growth of tumours over several days, to decades-long re-growth of corals in coral reefs that can extend for hundreds of kilometres. Different mathematical models and statistical methods are used to diagnose, understand, and predict biphasic growth. Common approaches can lead to inaccurate predictions of future growth that may result in inappropriate management and intervention strategies being implemented. Here we develop a very general computationally efficient framework, based on profile likelihood analysis, for diagnosing, understanding, and predicting biphasic population growth. The two key components of the framework are: (i) an efficient method to form approximate confidence intervals for the change point of the growth dynamics and model parameters; and, (ii) parameter-wise profile predictions that systematically reveal the influence of individual model parameters on predictions. To illustrate our framework we explore real-world case studies across the life sciences.

## 1 Introduction

Quantifying population growth, whether it be of the total number of individuals in a group or of the total area covered by a species, motivated some of the earliest mathematical models [1]. The study of population growth is now significantly more sophisticated. Mathematical models are commonly employed to describe the growth of populations and the growth of individuals [2, 3, 4]. Here, we focus on populations and individuals that undergo two phases of growth, often called *biphasic growth*. Biphasic growth is prevalent across a wide range of applications in the life sciences, including: ecological applications, for example coral reef growth after a disturbance (Figure 1a) [14], growth of individual fish [15, 16] and turtles [17]; two-dimensional cell biology assays, for example proliferation and scratch-wound assays (Figure 1b) [5, 6]; three-dimensional cancer tumour spheroid cell biology experiments (Figure 1c) [7, 8]; drug release assays [9]; environmental decay of pathogens [10]; survival kinetics of bacteria [11]; pole elongation in mycobacteria [12]; and, bacterial interactions during food production [13]. Given the wide range of applications, a variety of mathematical and statistical methods have been developed in different disciplines to understand specific cases of biphasic population growth. Here, we develop a new computationally efficient general framework for diagnosing, understanding, and predicting biphasic population growth that is broadly applicable across the life sciences. The approach, based on profile likelihood analysis in combination with parameter-wise profile predictions, enhances the accuracy and reliability of previous methods. These improvements enable greater understanding of population growth dynamics and assist decision making.

**Figure 1:**
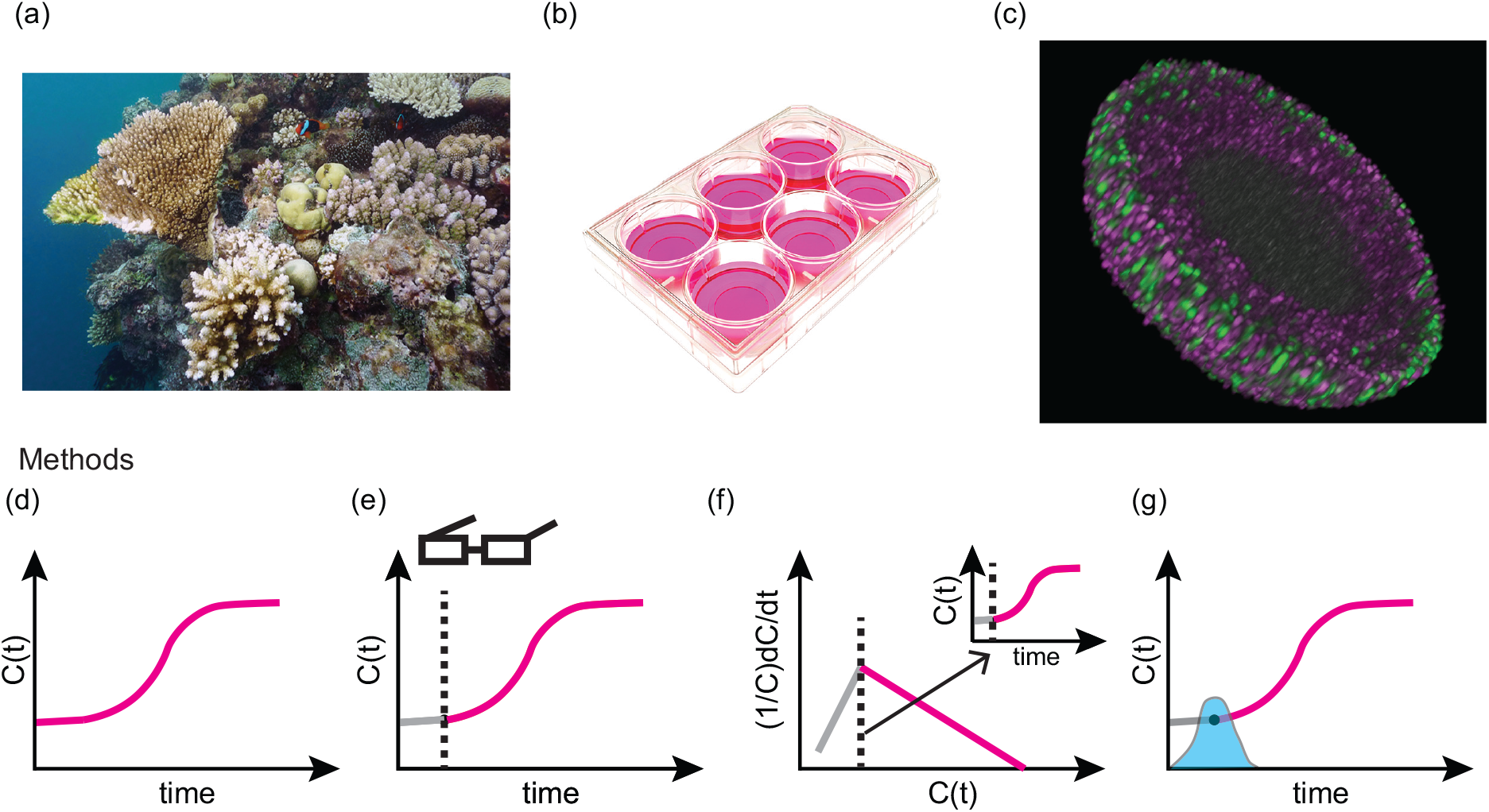
Case studies and methods for diagnosing, understanding, and predicting biphasic population growth in the life sciences. We explore biphasic population growth in: (a) coral reef growth after a disturbance [42]; (b) two-dimensional cell proliferation assays; and, (c) a three-dimensional cancer tumour spheroid experiment [7]. Existing methods to analyse biphasic population growth and the temporal evolution of a population *C*(*t*) range from: (d) overlooking the first phase (pink); (e) manually identifying the change point through visual inspection (first phase (grey), point estimate of change point (black-dashed), second phase (pink)); and, (f) analysing and identifying a change point in per capita growth rate data, 1*/C*(*t*) d*C*(*t*)*/*d*t* against *C*(*t*), and mapping this to a change point in time (first phase (grey), point estimate of change point (black-dashed), second phase (pink)). (g) In this study we form a profile likelihood for the change point (blue shaded) and each model parameter. Using the profile likelihoods we estimate approximate confidence intervals. To quantify and visualise how variations in a model parameter influence predictions we use parameter-wise profile predictions.

As biphasic population growth occurs across a wide range of applications and disciplines, different terminology is used to describe similar phenomena. A key term we refer to is the *change point* which is the time at which the growth dynamics switches from the first phase to the second phase. In cell biology and ecological applications, the first phase is sometimes referred to as a lag, delay, adaptation, or settling phase and the change point is sometimes referred to as the end of those respective phases or the start of the growth phase [5, 6, 14]. In the fisheries literature the change point is often referred to as the age of maturation, since the first phase represents growth as a juvenile and the second phase represents growth as an adult [15, 16]. Change point detection has a long history, with applications in signal analysis and econometrics [18, 19, 20], and standard tools have been developed in software such as MATLAB [21]. However, these standardised tools typically do not incorporate a mechanistic model. In contrast, mechanistic model-based approaches usually assume a specific model or do not provide a systematic statistical framework to assess uncertainty in change point estimates. Here we aim to bridge this gap by developing a general differential equation-based framework that does not rely on a specific model form while also providing systematic statistical uncertainty quantification.

Existing methods to analyse biphasic population growth vary in terms of simplicity, accuracy, and reliability. The simplest method to interpret biphasic population growth is overlook the two phases and analyse the experimental data with a single-phase model (Figure 1d) [14, 22, 23, 24]. Other approaches explicitly account for the existence of the two phases of growth and identify the change point manually through visual inspection (Figure 1e) [5, 7, 8]. More sophisticated methods involve seeking statistical point estimates of the change point. In econometrics this is sometimes referred to as a regression discontinuity study or two-segment regression with change point detection [18, 19]. Recent studies explore noisy per capita growth rate data to identify a change point in time (Figure 1f) [6, 14]. In another recent study examining the growth of individual fish [15], profile likelihood analysis has been used to form an approximate confidence interval for the change point (Figure 1g), albeit for a specific mathematical model.

When using differential equations to describe and interpret data one should consider whether model parameters are identifiable. Many studies focus on the formal question of *structural identifiability*, namely whether parameters of the mathematical model be uniquely identified given a set of continuous noise-free observations [25, 26, 27, 28]. Such analysis can be performed using software tools, such as `DAISY` [29] or `GenSSI` [30]. However, such tools focus on differential equations that are described by smooth functions and do not apply to biphasic growth models that are defined piecewise. Here, we focus on *practical identifiability*, namely whether given a finite set of noisy experimental data can we uniquely identify model parameters. Profile likelihood analysis is one approach to assess parameter identifiability [31, 32, 33, 34, 35, 36, 37]. We choose to base our framework on profile likelihood analysis for two key reasons: (i) computational efficiency [38]; and, (ii) to introduce parameter-wise profile predictions to quantify and visualise how variations in a model parameter influence predictions of population growth trajectories. Alternative approaches to assess parameter identifiability include Markov chain Monte Carlo techniques [39, 40, 41].

To illustrate our framework we explore four case studies across the life sciences: (i) coral reef re-growth after a disturbance; (ii) two different examples of two-dimensional cell proliferation assays; and, (iii) a three-dimensional cancer tumour spheroid experiment. In Section 2 we describe the various experimental and field-scale data sets. In Sections 3-5 we detail the mathematical model, techniques for parameter estimation, practical identifiability analysis and prediction intervals, including parameter-wise profile predictions. In Section 6 we apply our framework and in Section 7 we discuss insights that are gained by using this new framework.

## 2 Data

In this section we describe the data used in this study. Since we deal with two different proliferation assay experiments we present one of these cases, based on a bladder cancer cell line, in Supplementary Material F.

### 2.1 Coral reef growth after disturbance

Coral reef data analysed in this study are published in [14, 43] and are part of the Australian Institute of Marine Science’s Long Term Monitoring Program. The data describes the temporal evolution of the percentage coral cover following a major storm disturbance event (19 November 2008 to 18 September 2018) at Broomfield Island located within the Great Barrier Reef, Australia (Figure 2a).

**Figure 2:**
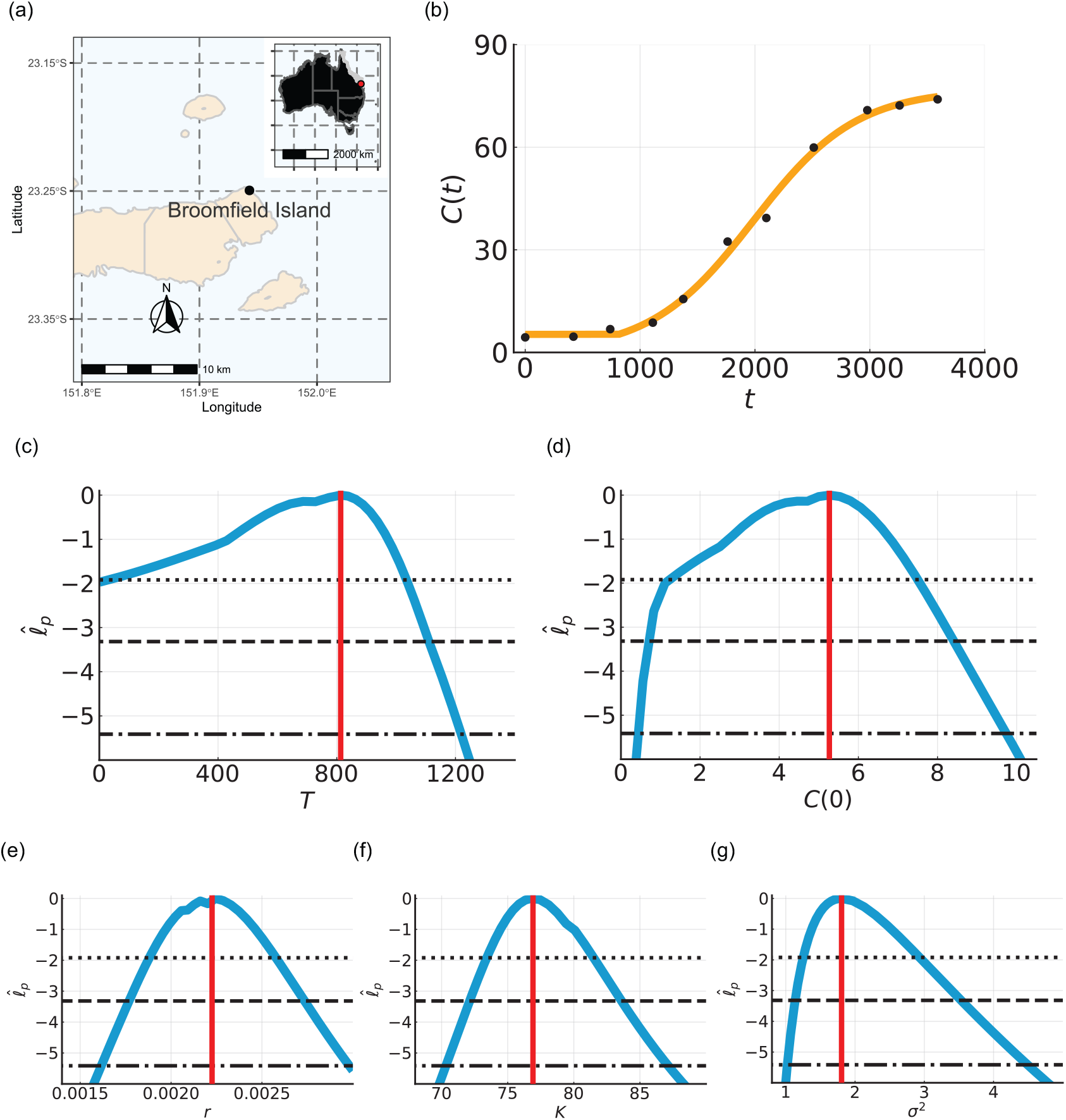
Biphasic coral reef growth after a disturbance. (a) The coral cover percentage, *C*(*t*) [%], was measured at Broomfield Island, Great Barrier Reef, Australia. (b) Comparison of the mathematical model simulated with the MLE (orange line) and field data (black circles) for the coral cover percentage, *C*(*t*) [%]. (c-g) Profile likelihoods for (c) *T* [days], (d) *C*(0) [%], (e) *r* [days^*−*1^], (f) *K* [%], and (g) *σ*^2^ [-] (blue) together with the MLE (vertical red line) and approximate 95% (dotted), 99% (dashed), and 99.9% (dash-dotted) confidence interval thresholds. The approximate 99.9% confidence intervals are: (c) *T* ∈ (0, 1220) [days], (d) *C*(0) ∈ (0.42, 9.74) [%], (e) *r* ∈ (0.016, 0.0030) [days^*−*1^], (f) *K* ∈ (70.3, 87.1) [%], and (g) *σ*^2^ ∈ (1.02, 4.50) [-].

### 2.2 Two-dimensional cell proliferation assay

This data set is obtained from an *in vitro* cell proliferation assay performed in [5]. A freshly prepared flask is placed in an incubator on a microscopic stage and the number of cells are observed as they divide to form a confluent monolayer. The experiment is performed on tissue culture plastic with NIH-3T3 fibroblast cells for 120 hours (5 days). Experimental measurements are normalised using the mean maximum cell density such that the normalised cell density ranges from zero to unity.

### 2.3 Three-dimensional cancer tumour spheroid experiment

This data set is obtained from tumour spheroid experiments we performed in [7, 8]. Human melanoma WM983b spheroids are formed with 5000 cells per well in a 96-well plate. Experiments are performed for 432 hours (18 days) and top-down area measurements of the spheroid are obtained using automated brightfield imaging and processing with the IncuCyte S3 live cell imaging system (Sartorius, Goettingen, Germany) (Supplementary Table S1). Images are captured every two hours for the first two days and then every six hours for the remainder of the experiment. In the first phase of dynamics the cells in the well migrate and adhere to form a shrinking spheroid. In the second phase of dynamics the spheroid grows as a solid mass. We quantify both phases by estimating the area enclosed by a projection of the spheroids, *A*, and assuming a spherical geometry convert these estimates into an equivalent radius 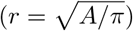. Results in the main manuscript deal with 0 *< t <* 120 [hours] and in Supplementary Material G we re-examine the data over a longer time interval 0 *< t <* 432 [hours].

## 3 Mathematical model

### 3.1 Process model

Let *C*(*t*) denote the variable of interest: for coral reef data this is coral cover percentage [14]; for two-dimensional cell proliferation assays this is the normalised cell density [5]; and for three-dimensional tumour spheroid experiments this is the radius of the tumour spheroid [7, 8]. To describe the population dynamics we prescribe a biphasic mathematical model,

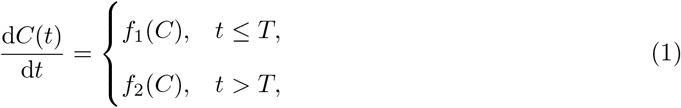

where *f*_1_(*C*) and *f*_2_(*C*) describe the time rate of change of *C*(*t*) before and after the change point, *t* = *T*, respectively. This framework is very general and can be used to describe several phenomena depending on how we specify *f*_1_(*C*) and *f*_2_(*C*). For example, if there is no growth or decay before *t* = *T* and logistic growth for *t > T*, we set *f*_1_(*C*) = 0 and *f*_2_(*C*) = *rC*(1 − *C/K*), where *r >* 0 is the growth rate and *K >* 0 is the long-time carrying capacity. For this application we have four unknown parameters, i.e. a vector (*r, K, C*(0), *T*), that we will estimate from data. For this particular choice of *f*_1_(*C*) and *f*_2_(*C*) we can solve the model exactly to give *C*(*t*) = *C*(0) for *t* ≤ *T* and *C*(*t*) = *KC*(0)*/* [*C*(0) + (*K* − *C*(0))exp(−*r*(*t* − *T*))] for *t > T*. Although, in principal, we can solve for *C*(*t*) exactly for certain choices of *f*_1_(*C*) and *f*_2_(*C*), all results presented in this work involve solving the mathematical model numerically using a second-order explicit Runge-Kutta method which means that we do not have to rely on integrating Equation (1) to obtain a closed-form solution.

### 3.2 Observation model

We assume that observed data 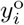 are measured at *I* discrete times, *t*_*i*_, for *i* = 1, 2, 3, …, *I*. We use a superscript ‘o’ to distinguish the noisy observed data from the model predictions. The model predictions are denoted by *y*_*i*_(*r, K, C*(0), *T*) = *C*(*t*_*i*_ | *r, K, C*(0), *T*). We collect the (noisy) data into a vector denoted by 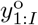. Similarly the process model solution is denoted by *y*_1:*I*_ (*r, K, C*(0), *T*) for the vector of grid point values and by *y*(*r, K, C*(0), *T*) for the full model trajectory over the time interval of interest. We estimate the process model parameter vector (*r, K, C*(0), *T*) by assuming that the observed data are noisy versions of the model solutions of the form 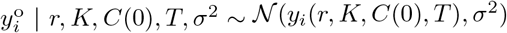. This means that we assume the observation error is additive and normally distributed with zero mean and constant variance *σ*^2^. Different error models could also easily be used within our likelihood-based framework. Here, the constant variance will be estimated along with the process model parameters.

## 4 Parameter estimation

We hence combine both the process model parameter vector (*r, K, C*(0), *T*) and the observation parameter *σ*^2^ into an overall vector parameter *θ* = (*r, K, C*(0), *T, σ*^2^). We can then consider scalar or vector sub-parameters as interest parameters defined as functions of the full vector parameter; for example *σ*^2^ = *σ*^2^(*θ*), where we use the same symbol for the function and its value. The process model solution is itself an interest parameter in this sense, and does not depend on the variance, i.e. *y*_*i*_(*θ*) = *y*_*i*_(*r, K, C*(0), *T, σ*^2^) = *y*_*i*_(*r, K, C*(0), *T*). Putting these elements together, we hence write our model for the data given the full parameter compactly as

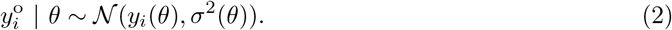

Taking a likelihood-based approach to parameter inference and uncertainty quantification, given a time series of observations together with our assumptions about the process and noise models, the log-likelihood function is

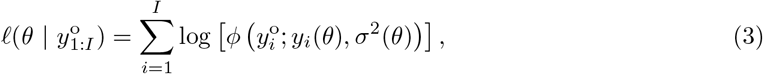

where *ϕ*(*x*; *µ, σ*^2^) denotes a Gaussian probability density function with mean *µ* and variance *σ*^2^. Maximum likelihood estimation (MLE) provides an estimate of *θ* that gives the best match (in the sense of highest likelihood) to the data. The MLE is given by

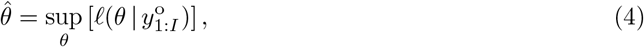

subject to bound constraints. The procedure for estimating 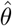 involves numerical maximisation of the log-likelihood which can be achieved using many different algorithms. In this work we find that a local optimisation routine from the open-source `NLopt` optimisation package in Julia performs well [44]. In particular, we use the Nelder-Mead optimisation routine within the `NLopt` with the default stopping criteria.

## 5 Practical identifiability analysis and profile predictions

We use a profile likelihood-based approach to explore practical identifiability by working with a normalised log-likelihood function

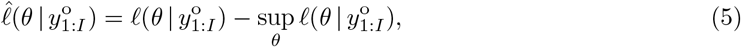

which we consider as a function of *θ* for a fixed set of data 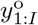. Note that normalising the log-likelihood means that we have 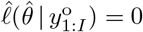.

### 5.1 Profile likelihood for interest parameters

Assuming the full parameter *θ* can be partitioned into an interest parameter *ψ* and nuisance parameter *λ*, where one or both of these may be vector-valued in general, we write *θ* = (*ψ, λ*). More generally we can consider an interest parameter as any well-defined function of the full parameter, *ψ* = *ψ*(*θ*), and leave the implied nuisance parameter implicit (that this always exists in the appropriate sense is implied by the results in [45]). For a set of data, 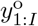, the profile log-likelihood for the interest parameter *ψ* given a partition (*ψ, λ*) is defined as [31, 46]

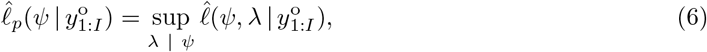

which indicates that *λ* is optimised out for each fixed value of *ψ*. This implicitly defines a function *λ*^*\**^(*ψ*) of optimal values of *λ* for each value of *ψ*. In the case of an interest parameter given as a general function of the full parameter, the profile (or induced) log-likelihood is defined in terms of the constrained optimisation problem [47, 48]

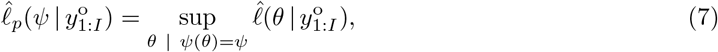

in which the ‘nuisance degrees of freedom’ in *θ*, after fixing *ψ*, are optimised out. As a concrete demonstration, consider the example in Sections 3-3.2 where we had *f*_1_(*C*) = 0 and *f*_2_(*C*) = *rC*(1 − *C/K*), and the full parameter vector is *θ* = (*r, K, C*(0), *T, σ*^2^). If we wish to profile the change point *T* then we have *ψ*(*θ*) = *T* and *λ*(*θ*) = (*r, K, C*(0), *σ*^2^) so that

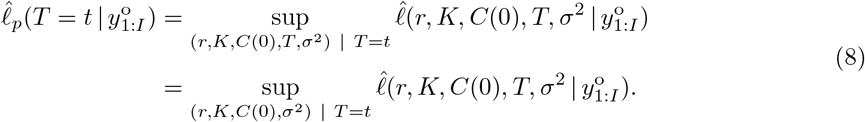

In all cases we implement this numerical optimisation using the same Nelder-Mead routine in `NLopt` that we use to estimate the MLE, 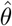 [44]. We define two uniformly-spaced meshes either side of the MLE in the interest parameter: (i) starting at the MLE to the lower bound of the interest parameter; and, (ii) starting from the MLE to the upper bound of the interest parameter. For all results in this work each mesh is formed by 40 points resulting in a total of 80 mesh points for each profile. For each mesh point to run the numerical optimisation we provide a starting estimate of the parameters. For the first mesh point closest to the MLE we set the starting estimate of *r, K, C*(0) and *σ*^2^ equal to their respective values in the MLE. We then seek the values of *r, K, C*(0) and *σ*^2^ that maximise 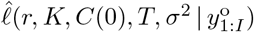. For the second mesh point closest to the MLE we use the estimate from the previous point as the starting estimate. For the starting estimate for all other mesh points we make a linear approximation using estimates at the previous two mesh points. The linear approximation holds provided the estimate remains within bounds. If it does not hold, we set the first guess as the previous estimate provided it remains within bounds and as the MLE otherwise. With these profiles, log-likelihood-based confidence intervals can be defined from the profile log-likelihood by an asymptotic approximation in terms of the chi-squared distribution that holds for sufficiently regular problems [31]. For example, 95%, 99% and 99.9% confidence intervals for a univariate (scalar) interest parameter correspond to a threshold profile log-likelihood value of −1.92, −3.32, and −5.41, respectively [49].

### 5.2 Predictive profile likelihood and parameter-wise profile predictions

Profile likelihoods for predictive quantities that are a (deterministic) function of the full parameter *θ* are defined in the same way as for any other function of the full parameter, as described in Section 5.1. For example, the full process model trajectory, *y*(*θ*), has an associated profile likelihood

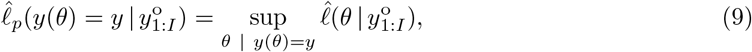

i.e. the profile likelihood value for a prediction is equal to the the maximum likelihood value across parameters consistent with that prediction. Here we give the profile prediction for the full (infinite-dimensional) model trajectory, which here is more straightforward in principle than general functional estimation problems as the variation is driven by a finite-dimensional parameter vector (and the constraint defined by the differential equation). However, this constraint may be more difficult to enforce in practice than the solution at a single time, and the literature typically focuses on a single-time prediction [33, 50]. In the special case that *y*(*θ*) is a one-to-one function, the above reduces to

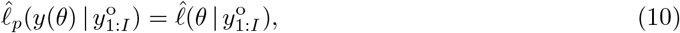

since the constraint *y*(*θ*) is uniquely invertible for *θ*. That is, profiling preserves the usual parameterisation invariance of the likelihood function under one-to-one transformations [47, 48]. However, profile predictions are still well-defined even without a one-to-one relationship between the parameters and model solution (i.e. in the absence of structural identifiability) [33] (see also [50, 51]).

Here we are also interested in some measure of the dependence of predictions on given target (interest) parameters. However, given a partition *θ* = (*ψ, λ*) and a function *q*(*θ*) of the full parameter, there is not in general a well-defined meaning of *q*(*ψ*), unless *q* is independent of *λ*. A natural approach then to exploring the dependence of a predictive function *q*(*ψ, λ*) of the full parameter on an interest parameter *ψ* is to consider its value along the corresponding profile curve, i.e. *q*(*ψ, λ*^*\**^(*ψ*)) where *λ*^*\**^(*ψ*) is the optimal value of the nuisance parameter for a given value of the interest parameter. We call these *parameter-wise profile predictions*, in contrast to the more standard predictive profile likelihood. In the simple case where *q* is independent of *λ* (or if *λ* is known) and is 1-1 in *ψ*, then this amounts to a re-parameterisation of the *ψ* profile likelihood. Hence, in this case, confidence intervals for *ψ* are directly transformed into confidence intervals for *q* (by transformation invariance of likelihood functions). The 1-1 requirement can be relaxed in the same way as for standard interest parameters but, in more complex cases with non-trivial dependence on the nuisance parameters, the transformation of confidence intervals for *ψ* into confidence intervals for the predictive quantity of interest will only be approximate and the precise statistical properties of these approximate prediction intervals are more difficult to establish (though can always be evaluated by simulation). In particular, if the predictive quantity of interest has weak or no dependence on the interest parameter being profiled and non-trivial dependence on the nuisance parameters, the associated predictive interval would be expected to have poor coverage. However, we can still use these parameter-wise intervals as an intuitive model diagnostic tool revealing the influence of an interest parameter on predictions. In contrast a standard predictive profile cannot reveal the individual influence of particular parameters. With these caveats in mind, we define the associated profile likelihood for *q*(*ψ, λ*^*\**^(*ψ*)) analogously to standard profile likelihood for an interest parameter, now starting from the profile likelihood for *ψ*:

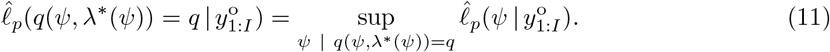

As with standard profile likelihood, this definition preserves parameterisation invariance under 1-1 transformations, i.e. if *q*(*ψ, λ*^*\**^(*ψ*)) is 1-1 in *ψ* then

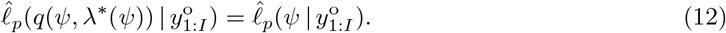

In addition to parameter-wise intervals, given a collection of individual intervals for the same quantity but based on different interest parameters, more conservative confidence intervals (relative to the individual intervals) for the predictions can be constructed by taking the union over all intervals. For example, given two intervals (or sets) 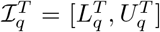 and 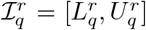 for a quantity *q* based on the profiles for *T* and *r*, respectively, we can form an interval (or set) 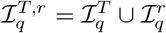 which has coverage at least as great as the individual intervals. In the case where the two intervals overlap, we have 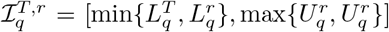. Again, the precise coverage properties of these intervals are difficult to establish, but such union intervals can provide an intuitive picture of overall variation in the predictive quantity.

## 6 Results and Discussion

Here we apply the general modelling framework that we present in Sections 3-5 to three case studies across the life sciences. We discuss a fourth case study, an additional two-dimensional cell proliferation assay that we perform with a bladder cancer cell line, in Supplementary Material F. These case studies cover a range of spatial and temporal scales, from microns and hours to kilometres and years, respectively.

### 6.1 Coral reef growth after disturbance

Recent modelling studies that examine the re-growth of coral reefs after some kind of disturbance (e.g. cyclone) have begun to explore the possibility that the re-growth involves a biphasic growth [14] whereas earlier studies have simply ignored this possibility [14, 22, 23]. Here, we explore measurements of the coral cover percentage, *C*(*t*) [%], of the reef at Broomfield Island, Great Barrier Reef, Australia (Figure 2a) [14]. In the first phase of growth *C*(*t*) remains approximately constant. In the second phase of growth *C*(*t*) is sigmoidal. Here we take the simplest approach and describe the second phase using the logistic growth model [23]. Therefore, we set *f*_1_(*C*) = 0 and *f*_2_(*C*) = *rC*(1 − *C/K*) in Equation (1), and seek estimates of five parameters, *θ* = (*T, C*(0), *r, K, σ*^2^).

Comparing the experimental data with the mathematical model simulated with the MLE, we observe very good agreement with small residuals that appear to be independent and identically distributed (Figure 2b). In terms of practical identifiability, the profile likelihood for *T* is wide, with approximate 99.9% confidence interval 0 ≤ *T* ≤ 1220 [days] and MLE at 813 [days]. This approximate confidence interval covers 1220 of 3590 days (34%) of the experimental data set. Therefore, it is unclear from these results whether there is no delay or a delay of three years. Understanding whether coral reef growth involves a delay is important for management and intervention strategies [14]. The profile likelihoods for *C*(0), *r, K*, and *σ*^2^ are relatively narrow and each well-formed around a single central peak suggesting these parameters are practically identifiable (Figures 2d-g).

To validate that the framework accurately estimates model parameters we repeat this analysis with two synthetic data sets based on the MLE of the coral reef experimental data (Supplementary Material B). The two synthetic data sets differ with respect to the value of *σ*^2^ used to generate the data. To generate the first data set we set *σ*^2^ equal to the MLE whereas for the second data set we reduce *σ*^2^. In both cases the framework accurately captures the known parameters used to generate the data. Reducing the variance suggests that the model parameters are structurally identifiable.

To improve our understanding of how each parameter influences mathematical modelling predictions we use parameter-wise profile predictions. We generate parameter-wise profile predictions for each of the five parameters and their union. A great advantage of using parameter-wise profile predictions is that we can identify the contribution of each of the parameters to predictions. First, we present the parameter-wise profile prediction for *T* (Figure 3a) and the difference between the parameter-wise profile prediction for *T*, 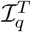, and the mathematical model simulated with the MLE, 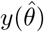, denoted 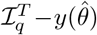 (Figure 3b). These results are very insightful since they show how uncertainty in each parameter affects different aspects of the predictions made using the model. For example, uncertainty in *T* is associated with a wider prediction interval at early time, but has very little impact upon the late time prediction interval (Figure 3a,b) which is intuitively reasonable since the late-time behaviour of the model is dictated by *K* rather than *T*. Similarly, we see that uncertainty in *C*(0) leads to a wide prediction interval at early time, but a smaller prediction interval at late time, which is also consistent with our understanding that *C*(0) plays in this model (Figure 3c-d). In contrast, uncertainty in *K* leads to a relatively wide prediction interval at late time, as expected, but a narrow prediction interval at early time (Figure 3g-h). As expected, *σ*^2^ provides zero contribution to the prediction of the mean due to the form of the error model. Given these parameter-wise prediction intervals we can then take the union of the parameter-wise profile predictions and understand how it is formed (Figure 3i-j). Inspecting the union of the parameter-wise profile predictions alone the contribution of each parameter to these differences is unclear.

**Figure 3:**
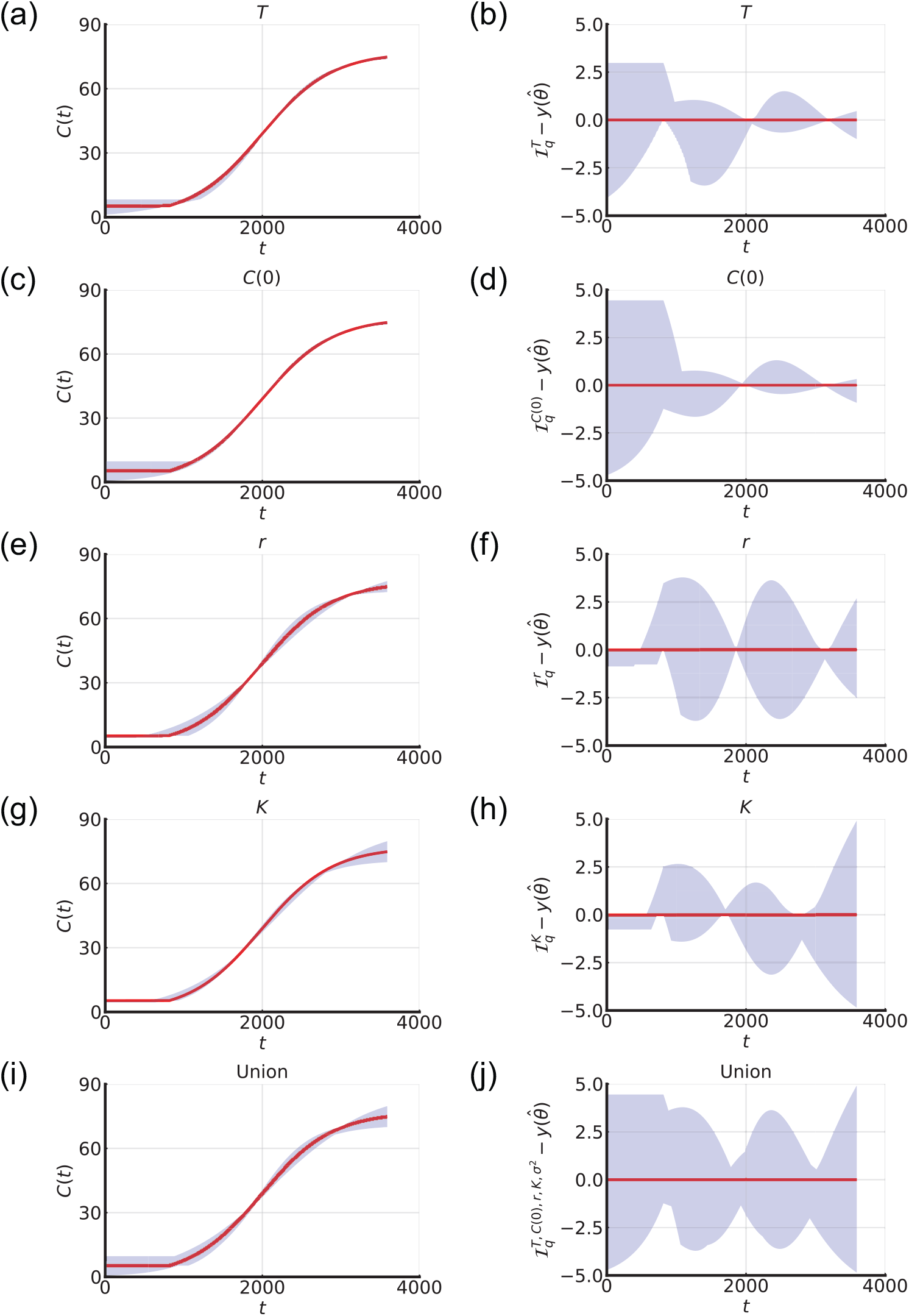
Parameter-wise profile predictions for coral reef growth. (a,c,e,g,i) Parameter-wise profile predictions for the mean (shaded) and the mathematical model simulated with the MLE (red). (b,d,f,h,j) Difference between parameter-wise profile predictions for the mean and the mathematical model simulated with the MLE. Results shown for (a,b) *T*, (c,d) *C*(0), (e,f) *r*, (g,h) *K*, and (i,j) the union of the parameter-wise profile predictions.

Many early studies of coral reef re-growth often ignore the possibility of biphasic growth (i.e. fixing *T* = 0) and do not allow for the possibility that *C*(0) can be estimated from the data (i.e. fixing *C*(0) equal to the first measurement) [14, 22, 23]. To demonstrate the impact of these more standard choices we repeat the analysis of this data under these assumptions (Figure 4). The mathematical model simulated with the MLE is fixed to capture the first data point (Figure 4a) but agreement to the other data points is considerably poorer in comparison to the biphasic model (Figure 2a). Furthermore, the residuals in this case are visually correlated, with systematic underestimation at early times and some overestimation at later times, violating statistical assumptions that the residuals are independent and identically distributed [23]. To compare the model where all parameters are estimated (Approach 1) with the model where we fix *T* = 0 and set *C*(0) equal to the first experimental measurement (Approach 2), we use the Akaike Information Criteria (AIC) [52]. The AIC is a standard tool for model selection studies and defined as 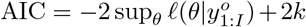 where *k* is the dimensionality of *θ* [53]. *When k* is the same for different models the AIC is a comparison of the maximum likelihood estimates and when *k* is different the model with more parameters is given a larger penalty. As the AIC is smaller for Approach 1 (54.5) than Approach 2 (69.5), this suggests that Approach 1 is more appropriate.

**Figure 4:**
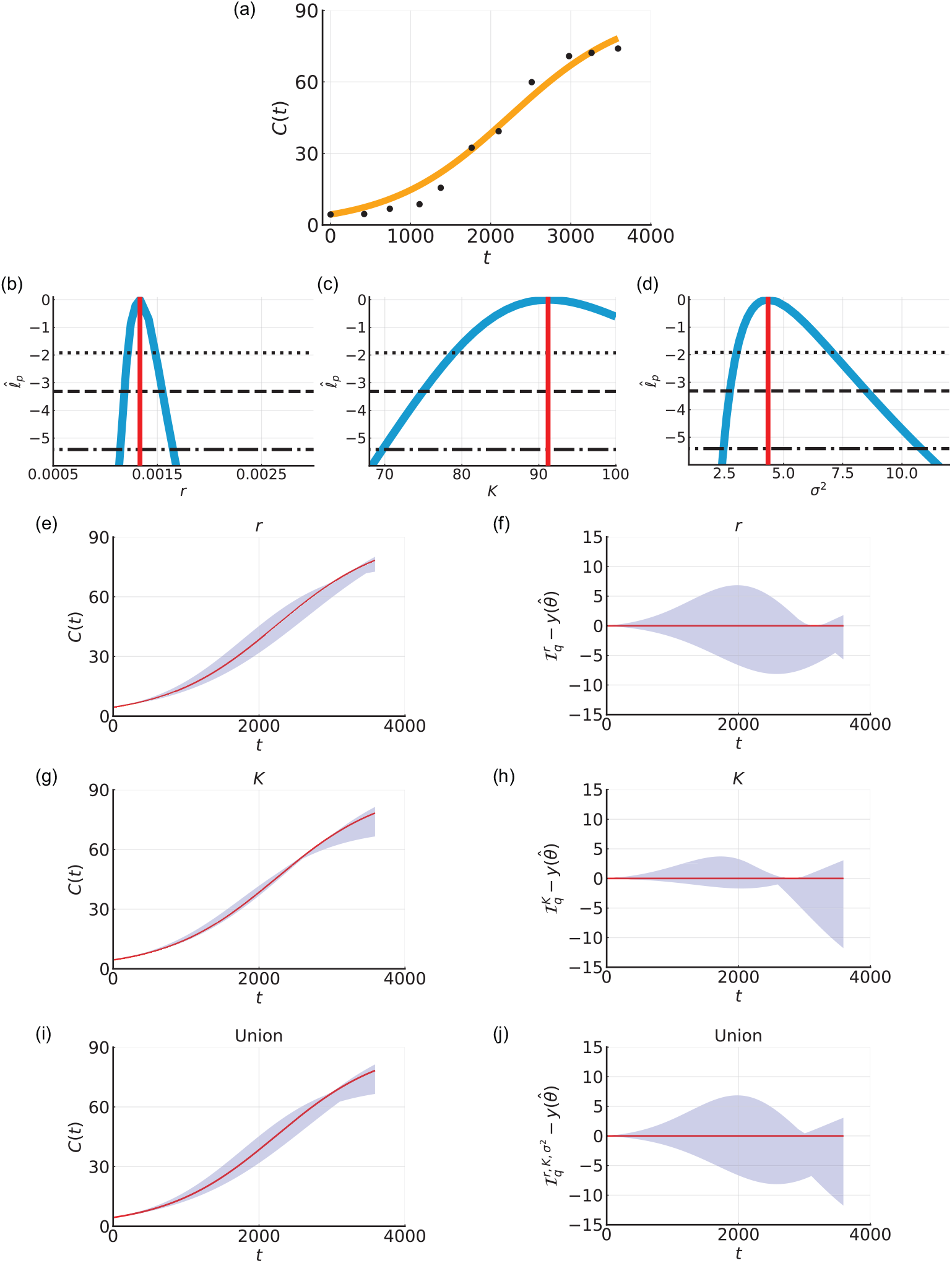
Biphasic coral reef growth after a disturbance: fixing *T* = 0 and setting *C*(0) equal to the first measurement. (a) Comparison of the mathematical model simulated with the MLE (orange line) and field data (black circles) for the coral cover percentage, *C*(*t*) [%], measured at Broomfield Island, Great Barrier Reef, Australia. (b-d) Profile likelihoods for (b) *r* [days^*−*1^], (c) *K* [%], and (d) *σ*^2^ [-] (blue) together with the MLE (vertical red line) and approximate 95% (dotted), 99% (dashed), and 99.9% (dash-dotted) confidence interval thresholds. The approximate 99.9% confidence intervals are: (b) *r* ∈ (0.0011, 0.0017) [days^*−*1^], (c) *K* ∈ (69.7, 100.00) [%], and (d) *σ*^2^ ∈ (2.46, 10.81) [-]. (e,g,i) Parameter-wise profile predictions for the mean (shaded) and the mathematical model simulated with the MLE (red). (f,h,j) Difference between parameter-wise profile predictions for the mean and the mathematical model simulated with the MLE. Results shown for (e,f) *r*, (g,h) *K*, and (i,j) the union of the parameter-wise profile predictions.

The AIC provides a single numerical value to compare mathematical models. Here, we provide further insights by comparing the profile likelihoods [23]. Profile likelihoods and approximate confidence intervals for parameters of the single-phase model are different to the corresponding profile likelihoods of the biphasic model. Specifically, estimates of *r* are smaller in the single-phase model than the biphasic model (Figures 2e, 4b). Furthermore, the approximate confidence interval for *K* is much larger for the single-phase model (Figures 2f, 4c). Such differences in parameter estimates could have major impacts on intervention and management strategies. For example, the single-phase model suggests it is likely that coral cover will eventually reach 100% (*K* = 100%), whereas *K* = 100% is a very unlikely prediction from the biphasic model. Results for a single-phase model (i.e. *T* = 0), without fixing *C*(0), are shown in Supplementary Material C.

### 6.2 Two-dimensional cell proliferation assay

Inspecting the time evolution of the normalised cell density, *C*(*t*) ∈ [0, 1] [-], in two-dimensional cell proliferation assays we observe biphasic population growth (Figure 5a). In the first phase of growth *C*(*t*) remains approximately constant. In the second phase of growth *C*(*t*) is sigmoidal. As before, we take the simplest approach and describe the second phase using the logistic growth model. Therefore, we set *f*_1_(*C*) = 0 and *f*_2_(*C*) = *rC*(1 − *C*) in Equation (1). We now seek estimates of four parameters, *θ* = (*T, C*(0), *r, σ*^2^).

**Figure 5:**
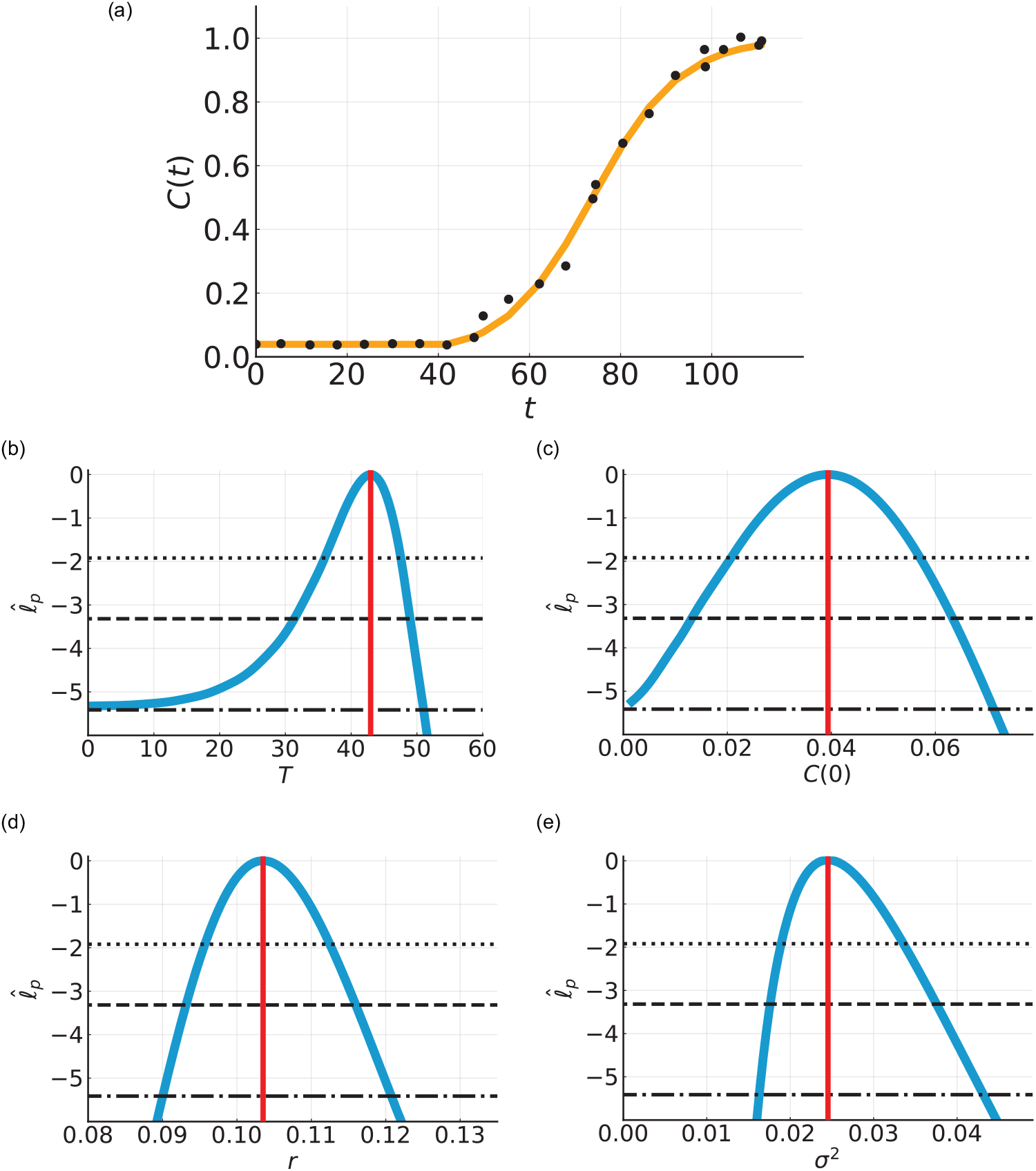
Biphasic population growth in a two-dimensional cell proliferation assay. (a) Comparison of the mathematical model simulated with the MLE (orange line) and experimental data (black circles) for the normalised cell density, *C*(*t*) [-]. (b-e) Profile likelihoods for (b) *T* [hours], (c) *C*(0) [-], (d) *r* [hours^*−*1^], and (e) *σ*^2^ [-] (blue) together with the MLE (vertical red line) and approximate 95% (dotted), 99% (dashed), and 99.9% (dash-dotted) confidence interval thresholds. The approximate 99.9% confidence intervals are: (b) *T* ∈ (0.0, 51.0) [hours], (c) *C*(0) ∈ (0.001, 0.071) [-], (d) *r* ∈ (0.090, 0.121) [hours^*−*1^], and (e) *σ*^2^ ∈ (0.016, 0.043) [-].

Comparing the experimental data with the mathematical model simulated with the MLE, we observe very good agreement with small visually uncorrelated residuals (Figure 5a). The profile likelihood for *T* is well-formed around a single central peak suggesting that *T* is practically identifiable to a 99% approximate confidence interval threshold (Figure 5b). However, the approximate 99.9% confidence interval is wider, 0 *< T <* 51 [hours], and the MLE is 43 [hours]. Previous analysis of this data set used visual inspection to estimate *T* = 40 [hours]. The approach we use here is more objective and reproducible and consequently more reliable and accurate than the previous method. Further, our approach provides an approximate confidence interval rather than a point estimate. Profile likelihoods for the three other parameters, *C*(0), *r*, and *σ*^2^, suggest they are practically identifiable (Figures 5c,d,e). Parameter-wise profile predictions reveal the influence of individual model parameters on predictions (Figure 6). Similar results are obtained for the fourth case study, a different cell proliferation assay experiment that we perform with a bladder cancer cell line and larger initial density (Supplementary Material F).

**Figure 6:**
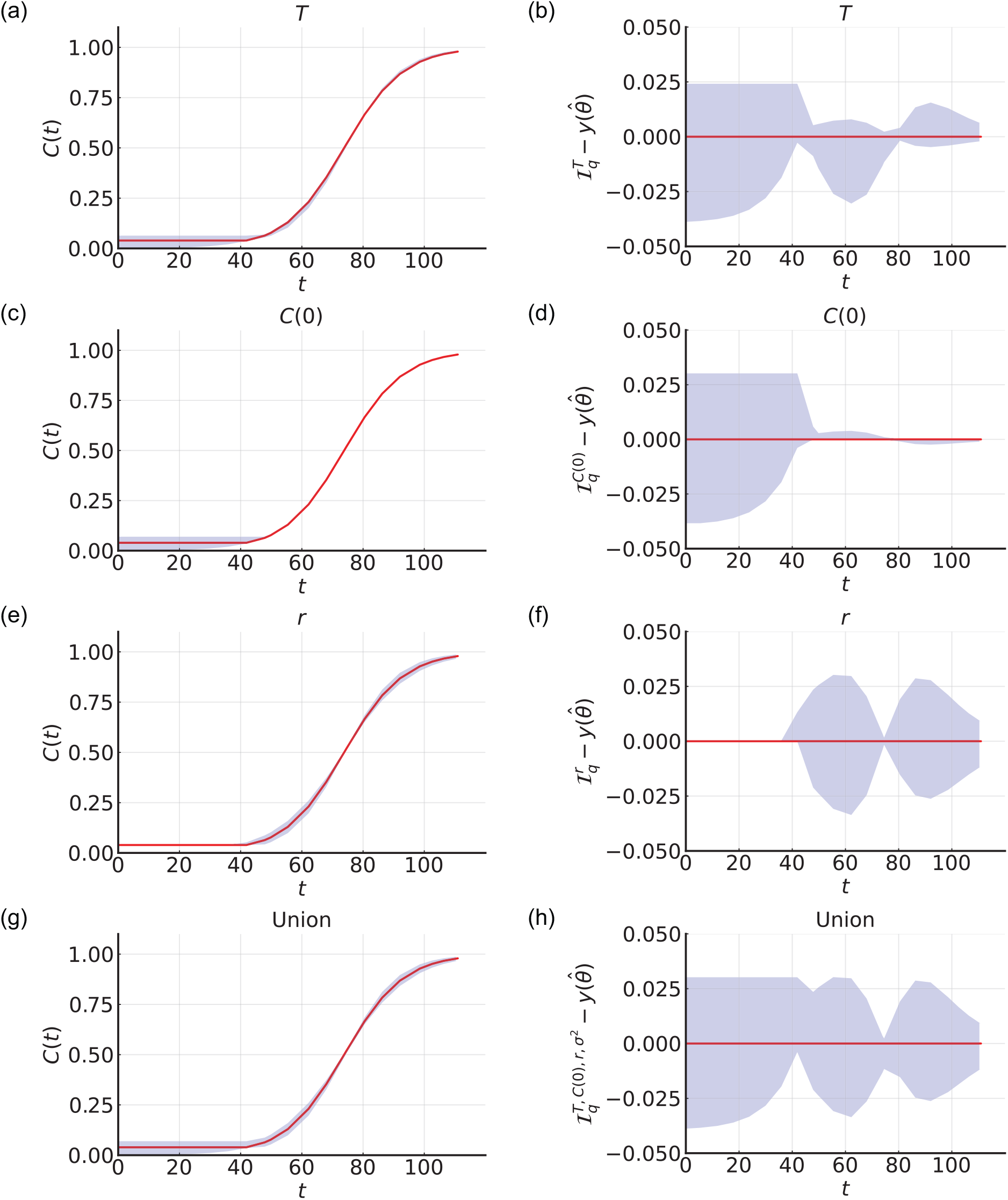
Parameter-wise profile predictions for two-dimensional cell proliferation assay. (a,c,e,g) Parameter-wise profile predictions for the mean (shaded) and the mathematical model simulated with the MLE (red). (b,d,f,h) Difference between parameter-wise profile predictions for the mean and the mathematical model simulated with the MLE. Results shown for (a,b) *T*, (c,d) *C*(0), (e,f) *r*, and (g,h) the union of the parameter-wise profile predictions.

### 6.3 Three-dimensional cancer tumour spheroid experiment

Three-dimensional tumour spheroid experiments performed in [7, 8] are initiated by placing a certain number of tumour cells into different wells of a tissue culture plate. The overall process of spheroid formation and growth involves two phases: in phase (i) the cells migrate and adhere to form a shrinking spheroid; and, in phase (ii) the newly formed spheroid grows as compact solid mass increases (Figures 1b, 7a-e). Over the entire experimental duration 0 *< t <* 432 [hours], *R*(*t*) increases to a long-time maximum radius, ℛ_2_ [µm] (Supplementary Material G). Here to illustrate the early-time biphasic behaviour we focus on 0 *< t <* 120 [hours].

Many models could be chosen to describe and analyse how *R*(*t*) evolves in time. Model selection has been well-studied for the second phase of growth [54, 55, 56], however, the first phase where the spheroid forms is rarely studied. Here we take a minimal approach and assume both phases can be described by distinct logistic growth models, giving

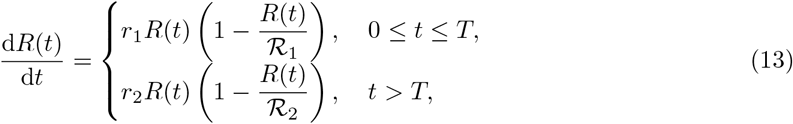

where *r*_1_ and *r*_2_ are the growth rates in the the first and second phase, respectively, and ℛ_1_ and ℛ_2_ are the associated limiting radii in each phase. Overall, we have seven parameters to estimate *θ* = (*T, R*(0), *r*_1_, *r*_2_, ℛ_1_, ℛ_2_, *σ*^2^). Using the logistic growth model to simulate growth of cell populations where the density is less than the long-time carrying capacity density is extremely common [4, 24, 54, 55, 56]. In contrast, using logistic growth where the dependent variable is greater than the long-time carrying capacity, as we do here to describe the first phase of spheroid formation, is quite unusual [57]. However, we find that this approach provides a good description of our experimental observations using a very familiar mathematical model.

Comparing the experimental data with the mathematical model simulated with the MLE, we observe very good agreement (Figure 7e). The profile likelihood for *T* is well-formed around a single central peak suggesting that *T* is practically identifiable to the 99.9% approximate confidence interval threshold (Figure 2b). Profile likelihoods suggest five of the six other parameters *C*(0), *r*_1_, ℛ_1_, and *σ*^2^ are practically identifiable (Figures 7g,h,i,j,l). The profile likelihood for ℛ_2_ is well-formed around a single central peak and practically identifiable to a 95% approximate confidence interval threshold (Figure 7k). However the approximate 99% and 99.9% confidence intervals are wider suggesting these parameters are practically non-identifiable using this data set (Figure 2b). Increasing the experimental duration narrows the confidence intervals for *r*_2_ and ℛ_2_ suggesting they are practically identifiable with appropriate additional data (Figures S12-S13). Parameter-wise profile predictions reveal the influence of individual model parameters on predictions (Figure 8). Here our framework improves on previous methods that use visual inspection to identify the start of the second phase of growth for analysis [7, 8].

**Figure 7:**
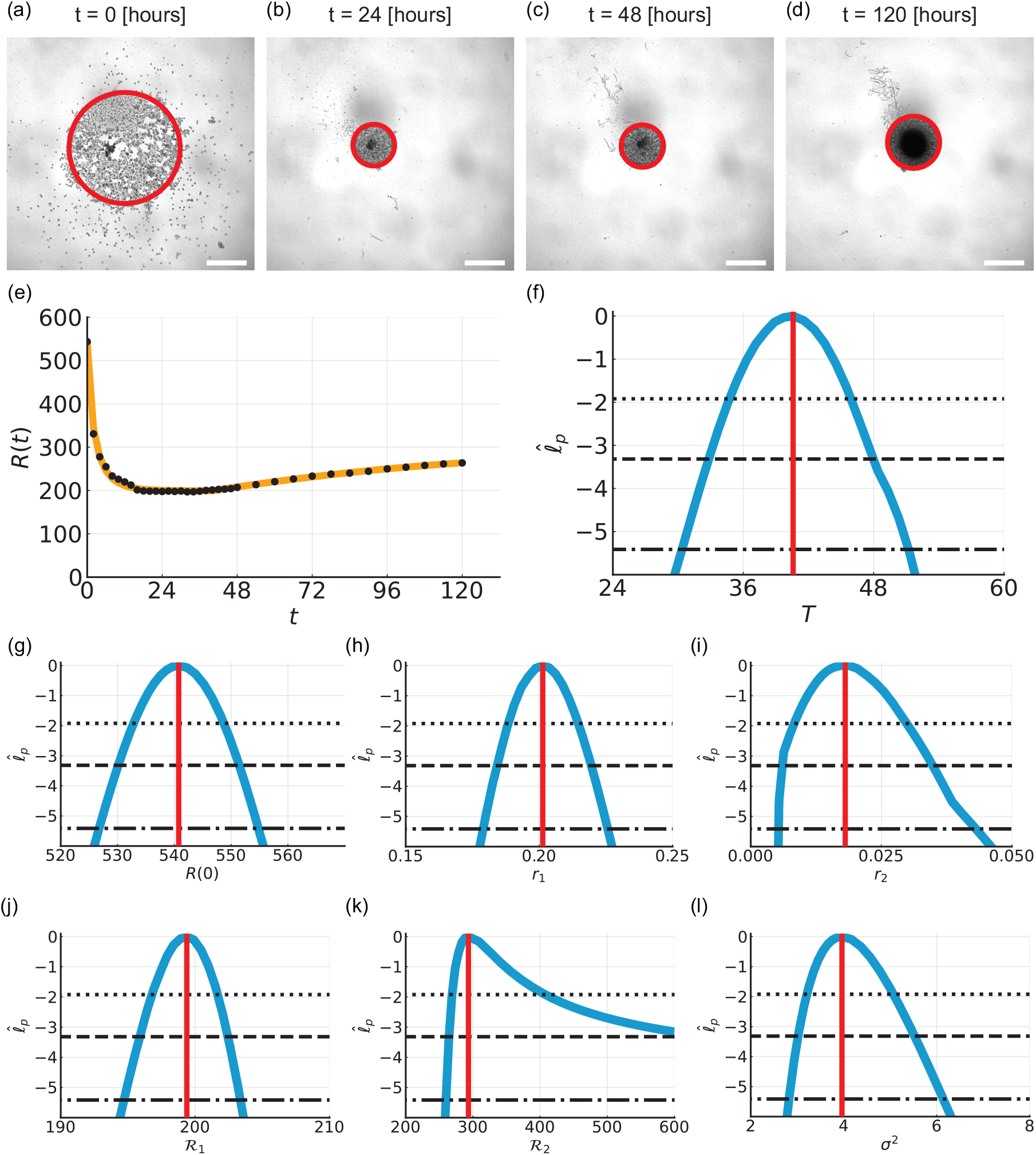
Biphasic population growth in three-dimensional tumour spheroid experiments. (a-d) Experimental images of spheroid experiments at (a) *t* = 0, (b) *t* = 24, (c) *t* = 48, and (d) *t* = 120 [hours]. Scale bar is 400 µm and red circle shows approximate equivalent area. At early times cells migrate and adhere to form spheroid. At later times the spheroid grows as a compact mass. (e) Comparison of the mathematical model simulated with the MLE (orange line) and experimental data (black circles) for the equivalent radius, *R*(*t*) [µm]. (f-l) Profile likelihoods for (f) *T* [hours], (g) *R*(0) [µm], (h) *r*_1_ [hours^*−*1^], (i) *r*_2_ [hours^*−*1^], (j) ℛ_1_ [µm], (k) ℛ_2_ [µm], and (l) *σ*^2^ [-] (blue) together with the MLE (vertical red line) and approximate 95% (dotted), 99% (dashed), and 99.9% (dash-dotted) confidence interval thresholds. The approximate 99.9% confidence intervals are: (f) *T* ∈ (30.4, 51.2) [hours], (g) *R*(0) ∈ (526.8, 554.8) [µm], (h) *r*_1_ ∈ (0.179, 0.226) [hours^*−*1^], (i) *r*_2_ ∈ (0.005, 0.043) [hours^*−*1^], (j) ℛ_1_ ∈ (194.7, 203.4) [µm], (k) ℛ_2_ ∈ (259.6, 600.0) [µm], and (l) *σ*^2^ ∈ (2.82, 6.14) [-].

**Figure 8:**
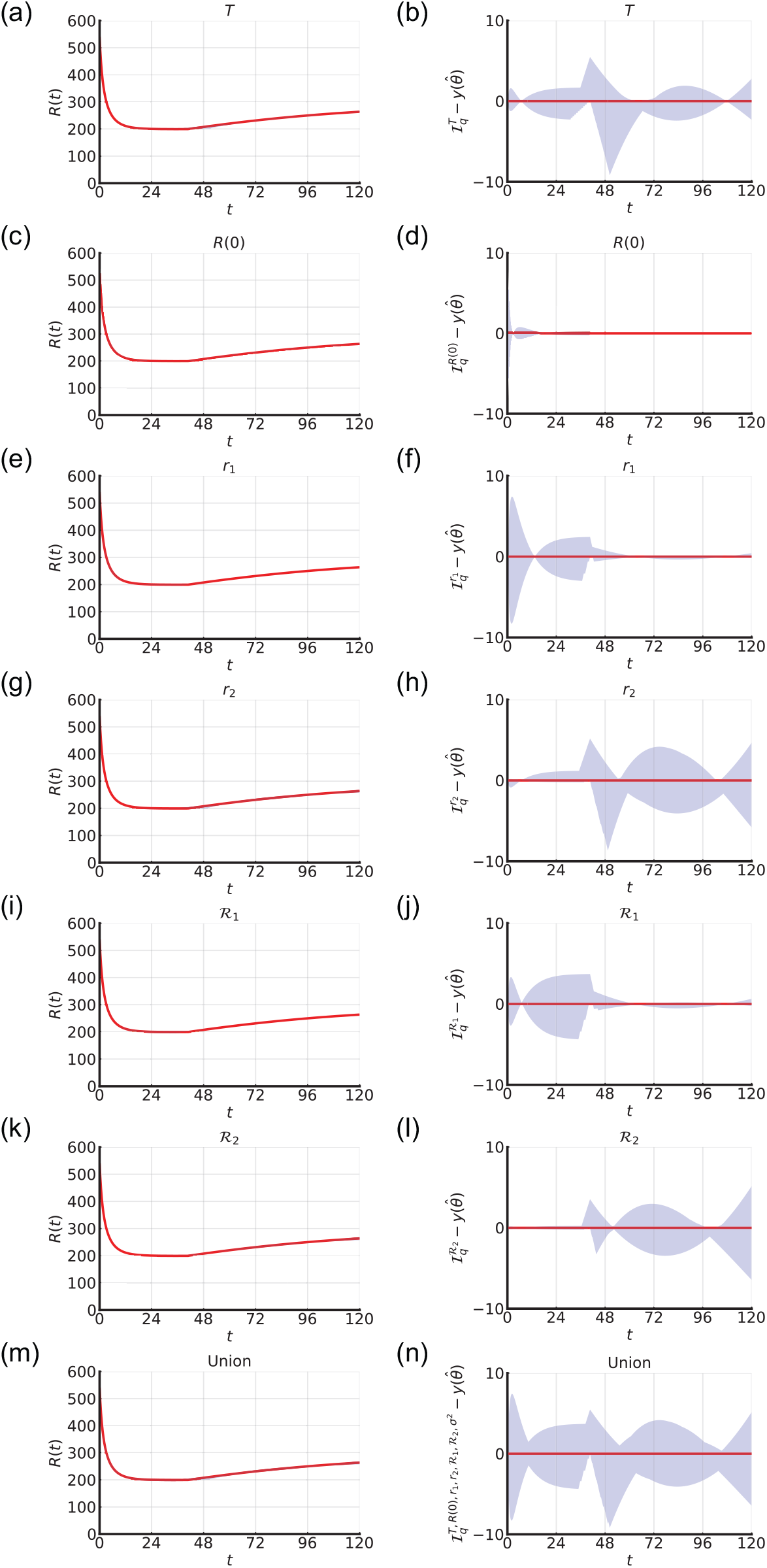
Parameter-wise profile predictions for three-dimensional cancer tumour spheroid experiment. (a,c,e,g,i,k,m) Parameter-wise profile predictions for the mean (shaded) and the mathematical model simulated with the MLE (red). (b,d,f,h,j,l,n) Difference between parameter-wise profile predictions for the mean and the mathematical model simulated with the MLE. Results shown for (a,b) *T*, (c,d) *R*(0), (e,f) *r*_1_, (g,h) *r*_2_, (i,j) ℛ_1_, (k,l) ℛ_2_, and (m,n) the union of the parameter-wise profile predictions.

## 7 Conclusion and Outlook

In this study we present a computationally efficient framework for diagnosing, understanding, and predicting biphasic population growth. Our framework involves two key components: (i) an efficient method to form approximate confidence intervals for the change point of the growth dynamics and model parameters; and, (ii) parameter-wise profile predictions that systematically reveal the influence of uncertainty for individual model parameters upon the model predictions. To demonstrate our framework we explore real-world case studies across the life sciences. This work builds on previous studies that focus on: single-phase models to describe biphasic growth; point-estimates of biphasic model parameters; and specific mathematical models and applications.

The ability to estimate the change point and model parameters in combination with parameter-wise profile predictions is powerful. For experimental design, parameter-wise profile predictions can inform when additional measurements should be taken to improve estimates of individual parameters. For the cell biology case studies, we provide accurate estimates of growth rates that can assist decision making in experiments, for example when to apply drug treatments [58]. For coral reef growth, we show that the framework can provide accurate parameter estimates and predictions that could aid management strategies. These case studies vary in terms of application and data quality, from sparse noisy data in coral reef studies to dense data collected in controlled experimental conditions in cell biology experiments. For all case studies the framework provides accurate parameter estimates and parameter-wise prediction intervals that lead to valuable insights.

Our work introduces parameter-wise prediction intervals in terms of the intuitive picture of variation in the predictive quantity. An open question from a theoretical point of view is how union intervals compare to standard profile predictive intervals for the same quantity. Because such parameter-wise (and union of parameter-wise) intervals are based on direct propagation of parameter uncertainties these are typically easier to compute than standard profile prediction intervals as the latter require enforcing constraints on the model outputs rather than inputs. On the other hand standard profile prediction intervals are more well-established theoretically.

This framework can be extended in many theoretical directions and to many applications. We take the simplest approach and use well-known logistic model to describe population dynamics. However, the framework is general and is well-suited to explore other models, for example Gompertz, generalised logistic, and Richard’s [23, 54, 55, 56]. Throughout we assume a Gaussian error model but different error models can easily be used within our likelihood-based framework. Furthermore, it is straightforward to extend the framework to explore growth dynamics that exhibit three or more growth phases, for example diauxic growth of bacteria [59]. One could also explore incorporating spatial effects by extending spatio-temporal single-phase partial-differential equation growth models [38, 60] to spatio-temporal biphasic growth models. Overall, this work lays the foundation for studies in biphasic population growth using differential equations, efficient change point and model parameter estimation, and parameter-wise prediction intervals.

## Data accessibility

Data and algorithms are available on GitHub.

## Author’s contributions

Conceptualization: RJM, OJM, MJS. Formal analysis: RJM, OJM, MJS. Funding acquisition: MJS. Methodology: All. Project administration: RJM, MJS. Resources: ARC, PBT, EDW. Software: RJM, OJM, MJS. Supervision: MJS. Visualisation: All. Writing - original draft: RJM, OJM, MJS. Writing - review and editing: All.

## Acknowledgements

We thank Dr Alexander P. Browning, Ms Gency Gunasingh, and Professor Nikolas K. Haaas for technical assistance and advice in the laboratory. MJS is supported by the Australian Research Council (DP200100177). DJW acknowledges support from the Centre for Data Science. EDW is supported by an award from the PA Research Foundation.

