## Supplementary Material for "Computationally efficient framework for diagnosing, understanding, and predicting biphasic population growth"

**Page No.**

A. Image processing for spheroid experiments 3

B. Validation with synthetic data 4

C. Coral reef growth: fixing $T = 0$ 9

D. Cell proliferation assay: fixing $T = 0$ 11

E. Cell proliferation assay: fixing $T = 0$ and $C(0)$ 13

F. Cell proliferation assay: bladder cancer cell line 15

G. Spheroid experiment: increased experimental duration 21

### A Image processing for spheroid experiments

Here we detail settings used in the IncuCyte S3 2020C Rev1 software for image processing. Settings from our previous study, focusing on spheroid growth after formation, are used for $t \geq 48$ [hours] [3]. For $t < 48$ [hours] the same settings do not provide accurate measurements of spheroid area or area covered by cells. Therefore, we use different settings (Table S1).

| Time [hours] | Channel | GCU threshold | Hole fill [µm$^2$] | Min area [µm$^2$] | Adjust size |
|---|---|---|---|---|---|
| 0 | green | 1 | $1.25 \times 10^7$ | $1 \times 10^5$ | 0 |
| 2, 4, 6 | green | 5 | $1 \times 10^7$ | - | 0 |
| 8, 10, 12 | green | 5 | $1 \times 10^7$ | - | -3 |
| 14 | green | 5 | - | - | -5 |
| 16-48 | red | - | $1 \times 10^7$ | - | 0 |
| $\geq$ 48 | red | - | - | - | 0 |

Table S1: Image processing IncuCyte settings. In this table a dash indicates the default setting. All measurements are obtained using the largest object area measure.

### B Validation with synthetic data

Here we generate synthetic data using a known mathematical model, known model parameters, and a known error model. We then show that our framework accurately recovers the known model parameters.

Focusing on the coral reef case study, we simulate the mathematical model with known model parameters ($r$, $K$, $C(0)$, $T$)=(0.00216, 77.36, 4.50, 701.92). Note these model parameters are the MLE of the coral reef experimental data set (Figure 2). For each time point in the coral reef experimental data set, we record the coral reef cover percentage from the mathematical model simulated with the known model parameters. Then for each recorded measurement from the simulated mathematical model we add noise by sampling from a normal distribution with zero mean and variance $\sigma^2$.

In the following we consider two values for $\sigma^2$, namely: (i) $\sigma^2 = 1.83$ corresponding to the MLE from the experimental data set (Figure S1); and (ii) a reduced variance, $\sigma^2 = 0.183$ (Figure S3). In both cases, the approximate confidence intervals capture the known parameters and the MLE's are close to the known parameters. As expected, when we reduce $\sigma^2$ the approximate confidence intervals narrow and the MLE's are found to be closer to the known parameter values. These results suggest that the mathematical model is structurally identifiable, which is useful since standard differential algebra techniques for assessing structurally identifiability do not apply when there is a discontinuity. Computing parameter-wise profile predictions for each of the five parameters and their union (Figures S2, S4), we show that the mathematical model simulated with the known parameters remains within the bounds of the union of the parameter-wise profile predictions (Figures S2j, S4j). Overall, these results validate that the framework accurately recovers the known parameters and dynamics used to generate the synthetic data.

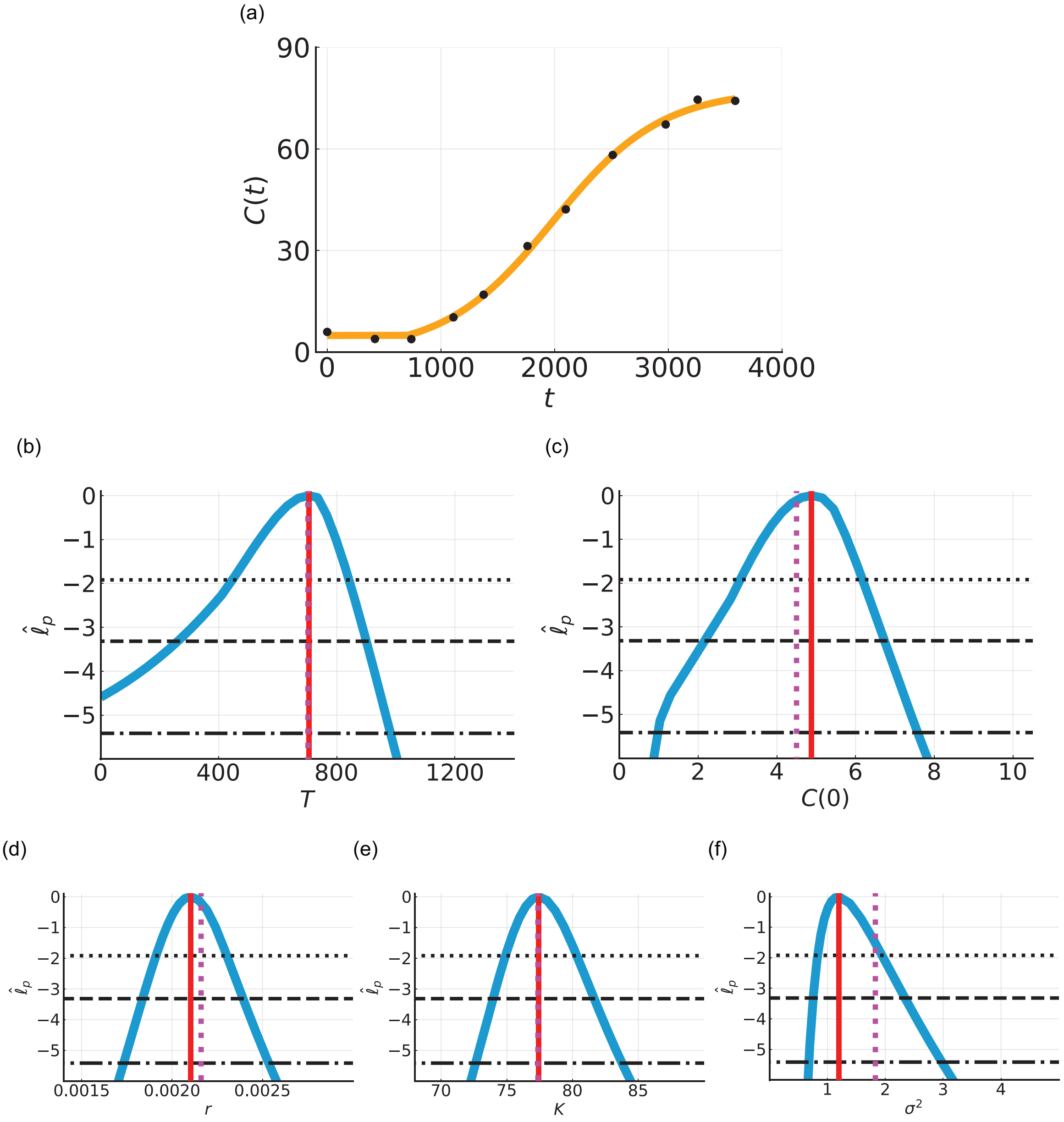


Figure S1: Synthetic data based on coral reef growth with known parameters ($r$, $K$, $C(0)$, $T$, $\sigma^2$)=(0.00216, 77.36, 4.50, 701.92, 1.823). (a) Comparison of the mathematical model simulated with the MLE (orange line) with synthetic data (black circles) for the coral cover percentage, $C(t)$ [%]. (b-f) Profile likelihoods for (c) $T$ [days], (d) $C(0)$ [%], (e) $r$ [days$^{-1}$], (f) $K$ [%], and (g) $\sigma^2$ [-] (blue) together with the known parameter value used to generate the data (vertical magenta line), MLE (vertical red line) and approximate 95% (dotted), 99% (dashed), and 99.9% (dash-dotted) confidence interval thresholds. The approximate 99.9% confidence intervals are: (b) $T \in (0, 954)$ [days], (c) $C(0) \in (0.86, 7.94)$ [%], (d) $r \in (0.0015, 0.0023)$ [days$^{-1}$], (e) $K \in (75.7, 90.0)$ [%], and (f) $\sigma^2 \in (0.81, 3.56)$ [-].

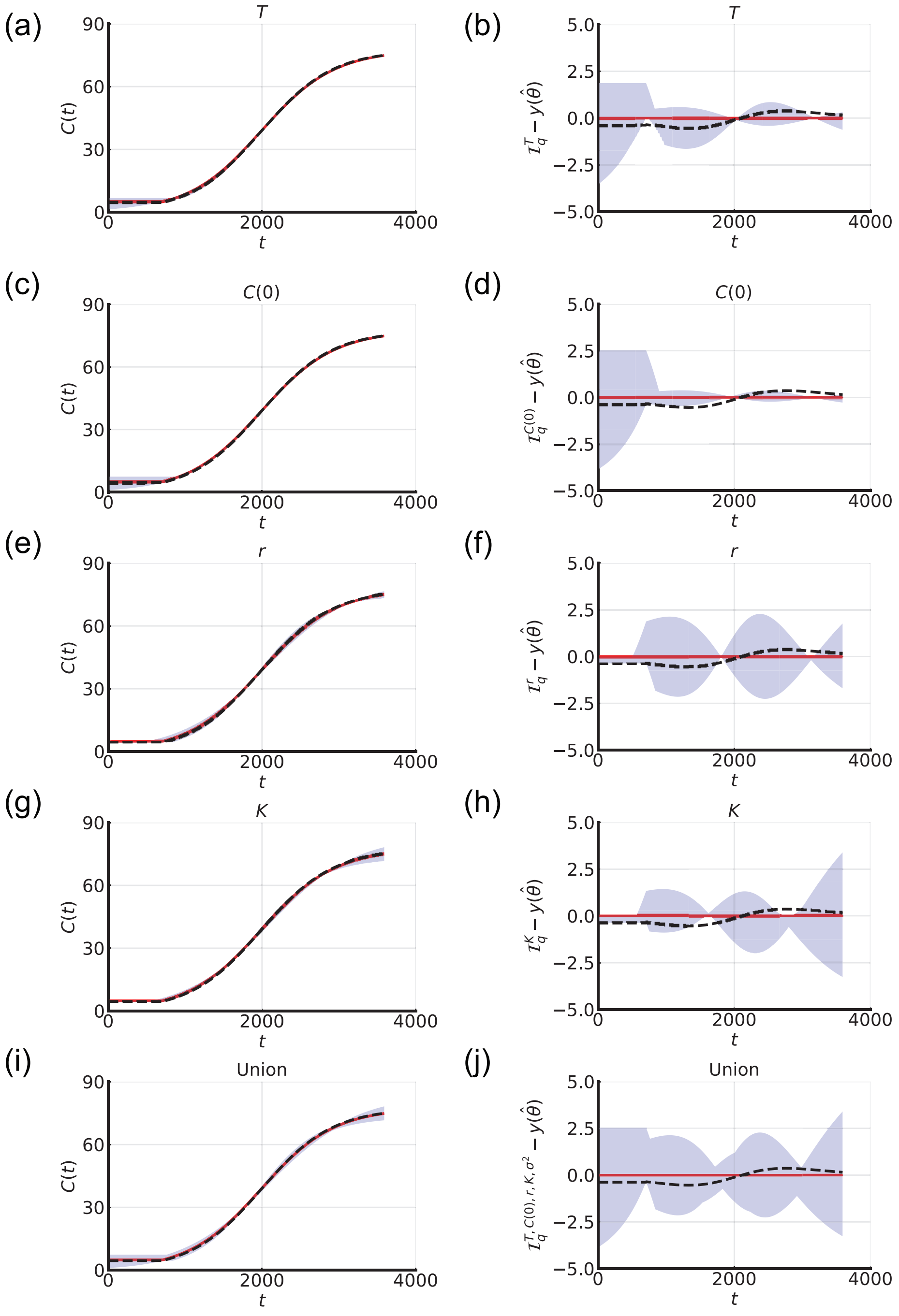


Figure S2: Parameter-wise profile predictions for synthetic data based on coral reef growth with known parameters ($r$, $K$, $C(0)$, $T$, $\sigma^2$)=(0.00216, 77.36, 4.50, 701.92, 1.823). (a,c,e,g,i) Parameter-wise profile predictions for the mean (shaded), the mathematical model simulated with the MLE (red), and the mathematical model simulated with the known parameters (black dashed). (b,d,f,h,j) Difference between parameter-wise profile predictions for the mean and the mathematical model simulated with the MLE (shaded). In (b,d,f,h,j) we also present the difference between the known solution and the mathematical model simulated with the MLE (black dotted). Results shown for (a,b) $T$, (c,d) $C(0)$, (e,f) $r$, (g,h) $K$, and (i,j) the union of the parameter-wise profile predictions.

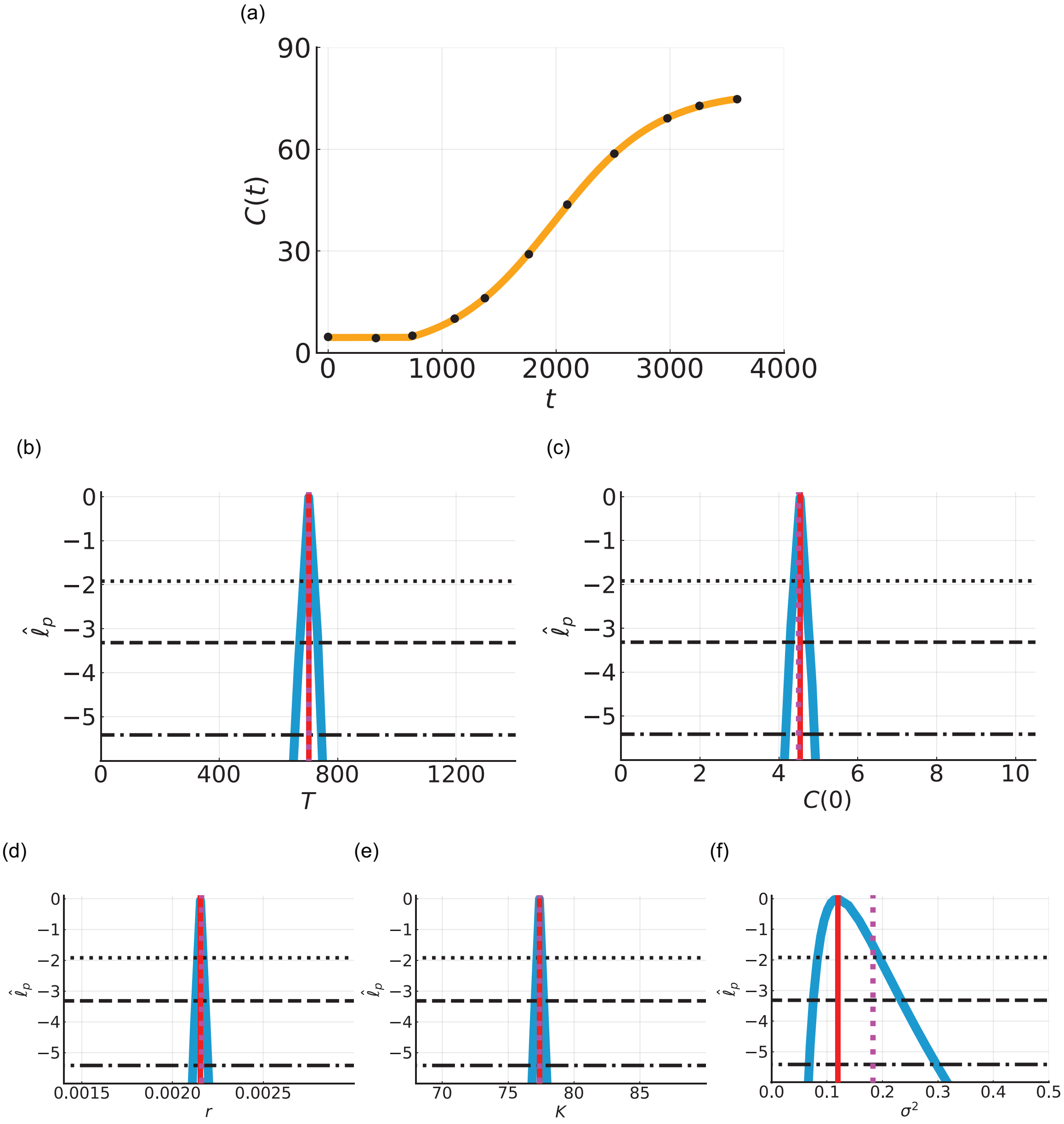


Figure S3: Synthetic data based on coral reef growth with known parameters and reduced variance ($r$, $K$, $C(0)$, $T$, $\sigma^2$)=(0.00216, 77.36, 4.50, 701.92, 0.1823). (a) Comparison of the mathematical model simulated with the MLE (orange line) with synthetic data (black circles) for the coral cover percentage, $C(t)$ [%]. (b-f) Profile likelihoods for (c) $T$ [days], (d) $C(0)$ [%], (e) $r$ [days$^{-1}$], (f) $K$ [%], and (g) $\sigma^2$ [-] (blue) together with the known parameter value used to generate the data (vertical magenta line), MLE (vertical red line) and approximate 95% (dotted), 99% (dashed), and 99.9% (dash-dotted) confidence interval thresholds. The approximate 99.9% confidence intervals are: (b) $T \in (654, 771)$ [days], (c) $C(0) \in (4.12, 5.06)$ [%], (d) $r \in (0.0021, 0.0022)$ [days$^{-1}$], (e) $K \in (76.5, 78.0)$ [%], and (f) $\sigma^2 \in (0.092, 0.402)$ [-].

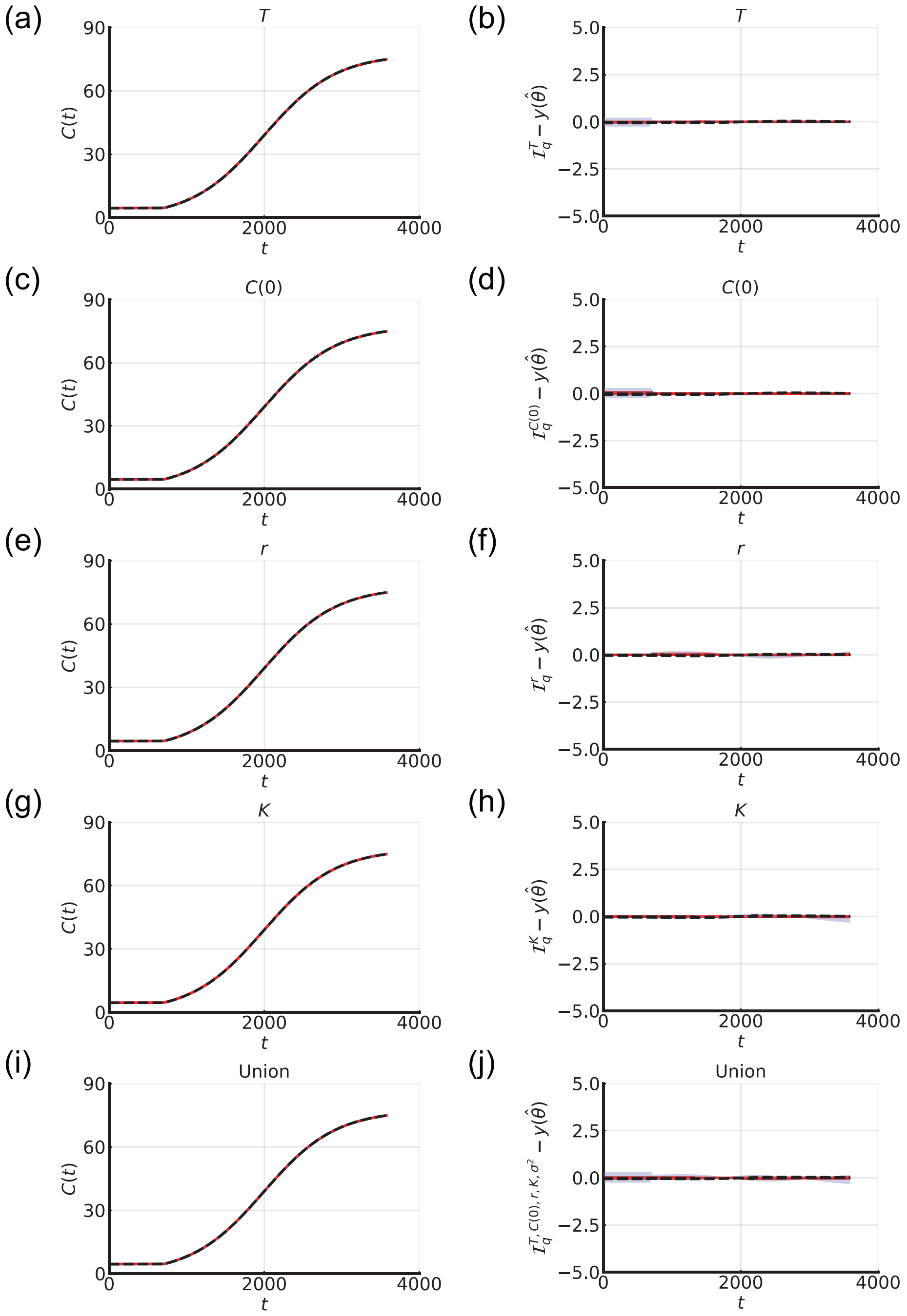


Figure S4: Parameter-wise profile predictions for synthetic data based on coral reef growth with known parameters and reduced variance ($r$, $K$, $C(0)$, $T$, $\sigma^2$)=(0.00216, 77.36, 4.50, 701.92, 0.1823). (a,c,e,g,i) Parameter-wise profile predictions for the mean (shaded), the mathematical model simulated with the MLE (red), and the mathematical model simulated with the known parameters (black dashed). (b,d,f,h,j) Difference between parameter-wise profile predictions for the mean and the mathematical model simulated with the MLE (shaded). In (b,d,f,h,j) we also present the difference between the known solution and the mathematical model simulated with the MLE (black dotted). Results shown for (a,b) $T$, (c,d) $C(0)$, (e,f) $r$, (g,h) $K$, and (i,j) the union of the parameter-wise profile predictions.

### C Coral reef growth: fixing $T = 0$

In main manuscript Section 6.1 we analyse coral reef growth data and estimate all parameters. Here, we analyse the same experimental data using a single-phase mathematical model (i.e $T = 0$) and estimate all other parameters ($C(0)$, $r$, $K$, $\sigma^2$) (Figure S5). These results suggest that the model does not accurately capture the first data point however other parameter estimates are similar to that of the biphasic model, albeit the approximate confidence intervals are generally wider.

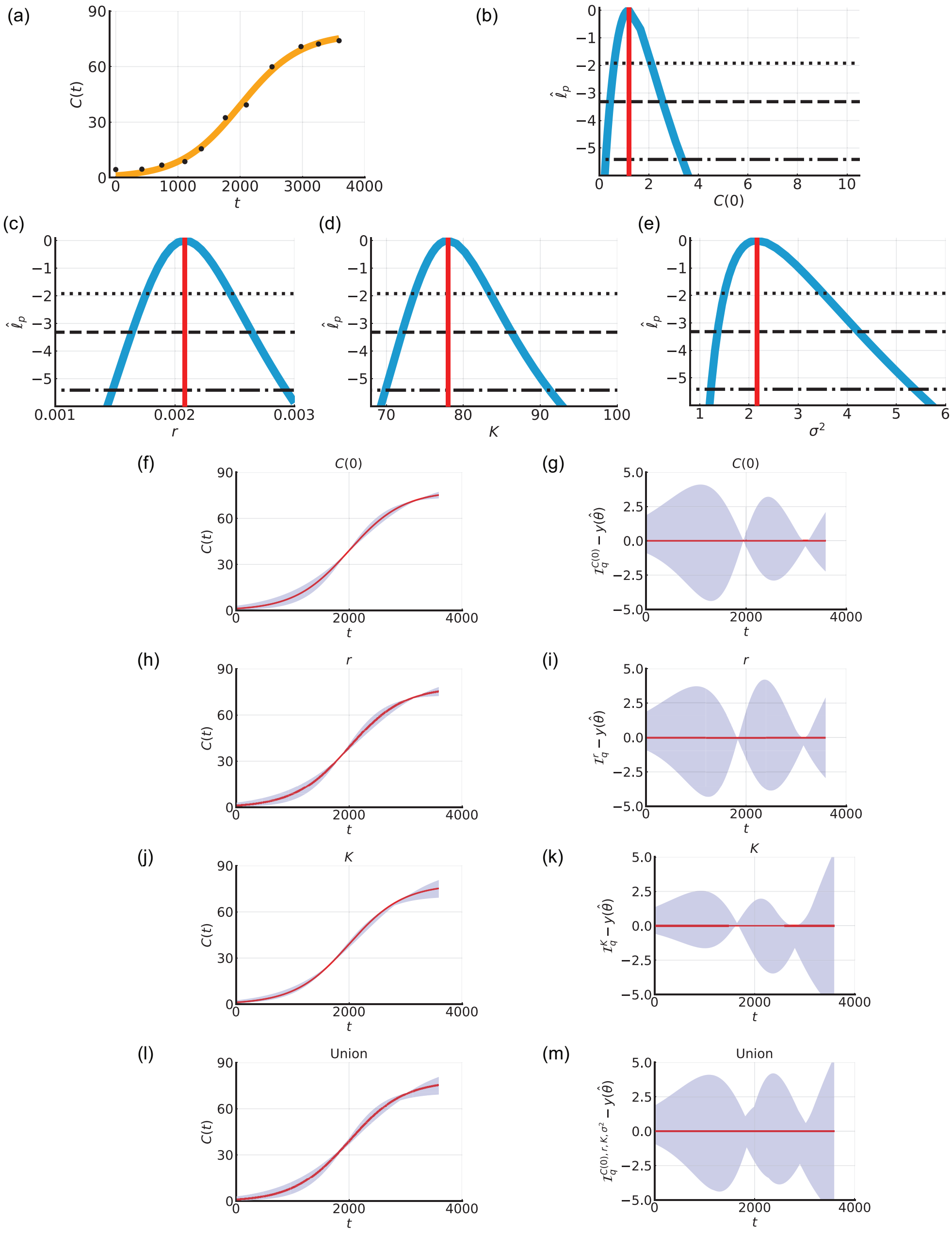


Figure S5: Biphasic coral reef growth after a disturbance: fixing $T = 0$. (a) Comparison of the mathematical model simulated with the MLE (orange line) and field data (black circles) for the coral cover percentage, $C(t)$ [%], measured at Broomfield Island, Great Barrier Reef, Australia. (b-e) Profile likelihoods for (b) $C(0)$ [%], (c) $r$ [days$^{-1}$], (d) $K$ [%], and (e) $\sigma^2$ [-] (blue) together with the MLE (vertical red line) and approximate 95% (dotted), 99% (dashed), and 99.9% (dash-dotted) confidence interval thresholds. The approximate 99.9% confidence intervals are: (b) $C(0) \in (0.25, 3.34)$ [%], (c) $r \in (0.0015, 0.0029)$ [days$^{-1}$], (d) $K \in (69.9, 91.2)$ [%], and (e) $\sigma^2 \in (1.22, 5.38)$ [-]. (f,h,j,l) Parameter-wise profile predictions for the mean (shaded) and the mathematical model simulated with the MLE (red). (g,i,k,m) Difference between parameter-wise profile predictions for the mean and the mathematical model simulated with the MLE. Results shown for (f,g) $C(0)$, (h,i) $r$, (j,k) $K$, and (l,m) the union of the parameter-wise profile predictions.

### D Cell proliferation assay: fixing $T = 0$

In main manuscript Section 6.2 we analyse the two-dimensional cell proliferation assay and estimate all parameters. Here, we analyse the same experiment with a single-phase mathematical model (i.e $T = 0$) and estimate all other parameters ($C(0)$, $r$, $\sigma^2$) (Figure S6) [4]. These results suggest that the model does not accurately capture the data points at early and intermediate times. Estimates for the other parameters are similar to the biphasic model, albeit the approximate confidence intervals are generally wider.

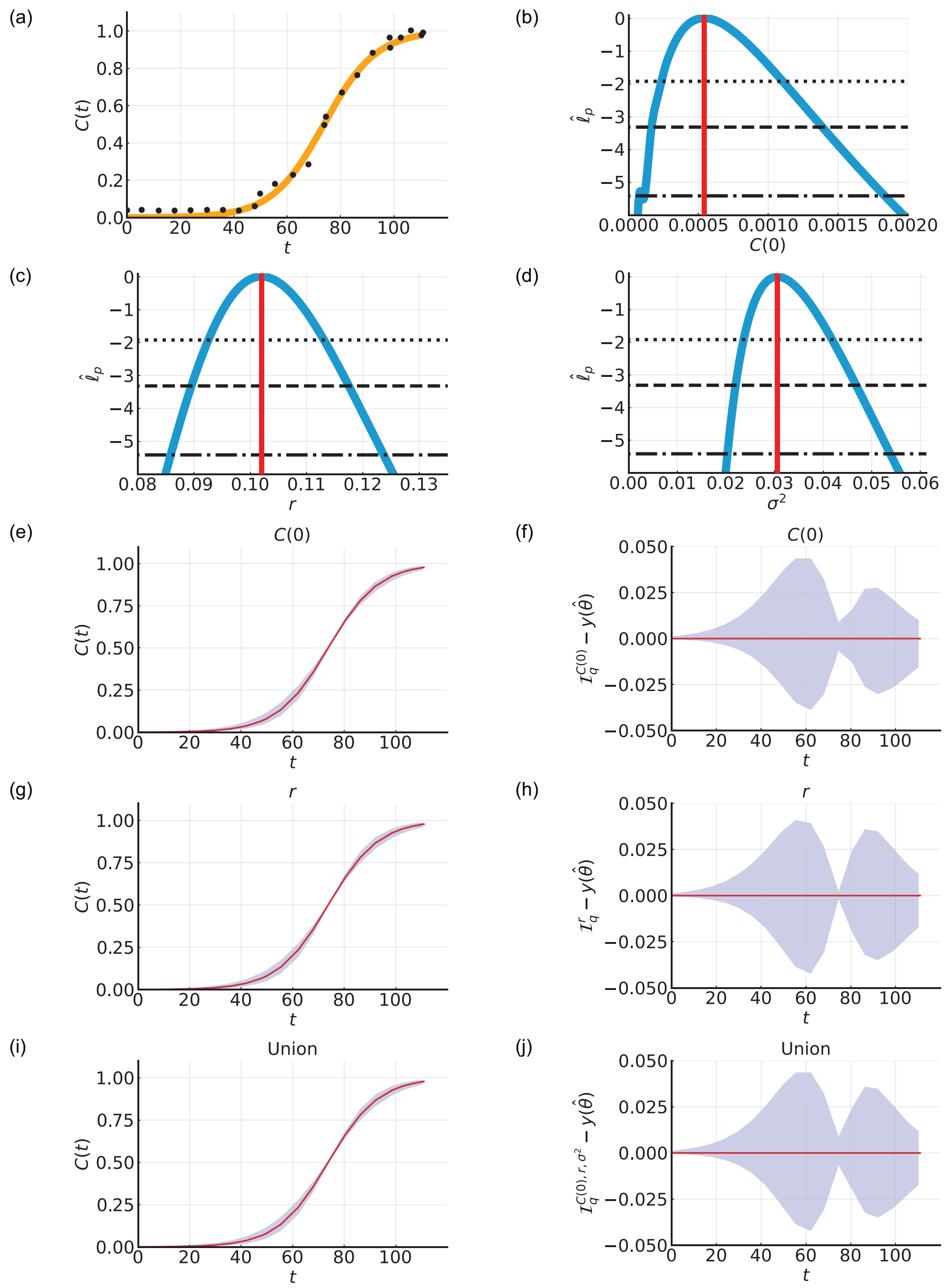


Figure S6: Biphasic population growth in a two-dimensional cell proliferation assay: fixing $T = 0$. (a) Comparison of the mathematical model simulated with the MLE (orange line) and experimental data (black circles) for the normalised cell density, $C(t)$ [-]. (b-d) Profile likelihoods for (b) $C(0)$ [-], (c) $r$ [hours$^{-1}$], and (d) $\sigma^2$ [-] (blue) together with the MLE (vertical red line) and approximate 95% (dotted), 99% (dashed), and 99.9% (dash-dotted) confidence interval thresholds. The approximate 99.9% confidence intervals are: (b) $C(0) \in (0.000, 0.002)$ [-], (c) $r \in (0.086, 0.124)$ [hours$^{-1}$], and (d) $\sigma^2 \in (0.020, 0.054)$ [-]. (e,g,i) Parameter-wise profile predictions for the mean (shaded) and the mathematical model simulated with the MLE (red). (f,h,j) Difference between parameter-wise profile predictions for the mean and the mathematical model simulated with the MLE. Results shown for (e,f) $C(0)$, (g,h) $r$, and (i,j) the union of the parameter-wise profile predictions.

### E Cell proliferation assay: fixing $T = 0$ and $C(0)$

In main manuscript Section 6.2 we analyse the two-dimensional cell proliferation assay and estimate all parameters. Here, we analyse the same experiment with a single-phase mathematical model (i.e $T = 0$), set $C(0)$ equal to the first experimental measurement, and estimate all other parameters ($r$, $\sigma^2$) (Figures S7) [4]. These results show that the mathematical model is fixed to capture the first data point. However, the match to the other data points is poor in comparison to the biphasic model. Furthermore, the residuals are correlated with systematic underestimation at early-intermediate times and overestimation at later times. Consequently, parameter estimates are not consistent with the biphasic model.

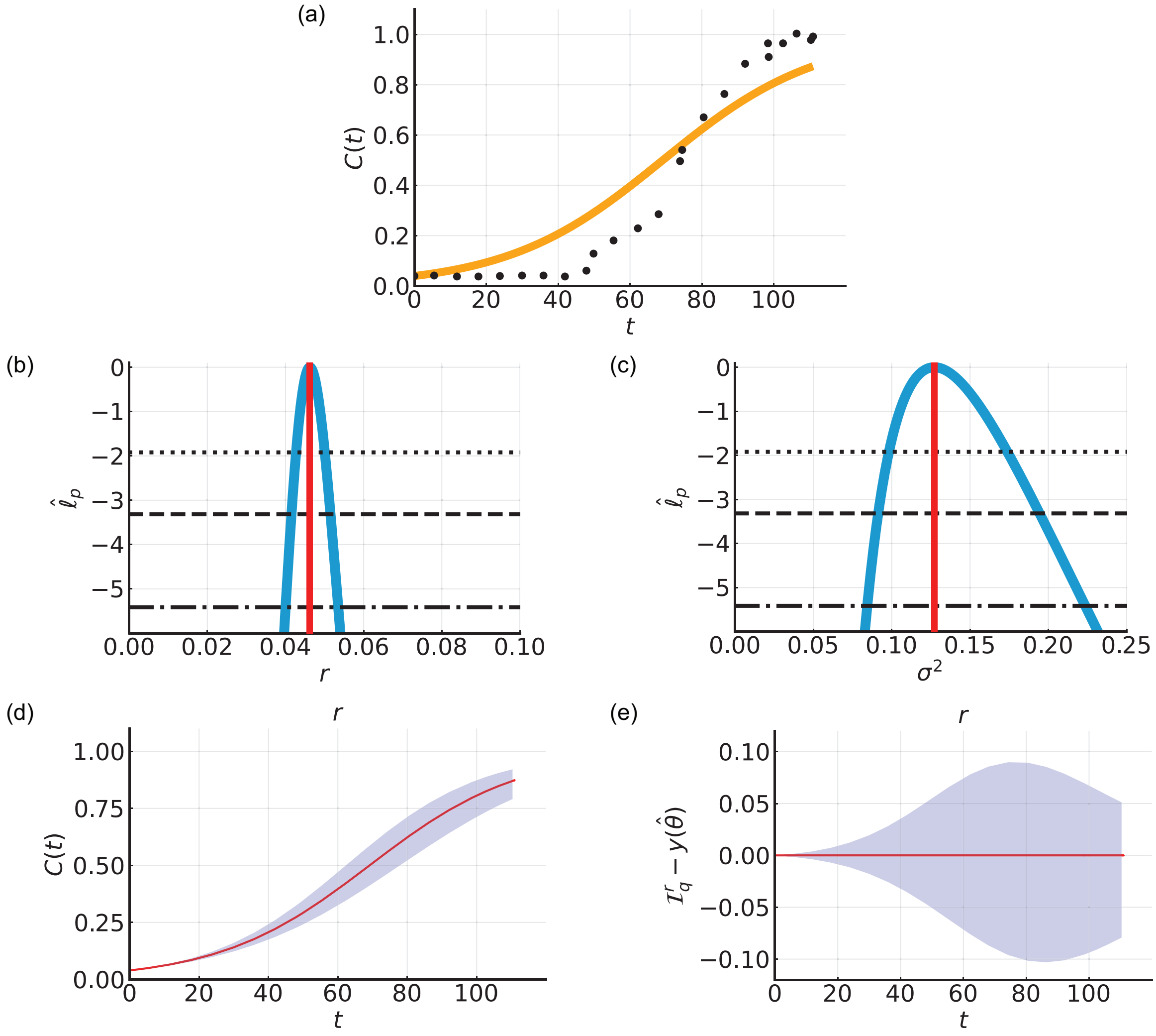


Figure S7: Biphasic population growth in a two-dimensional cell proliferation assay: fixing $T = 0$ and setting $C(0)$ equal to the first experimental measurement. (a) Comparison of the mathematical model simulated with the MLE (orange line) and experimental data (black circles) for the normalised cell density, $C(t)$ [-]. (b-c) Profile likelihoods for (b) $r$ [hours$^{-1}$], and (c) $\sigma^2$ [-] (blue) together with the MLE (vertical red line) and approximate 95% (dotted), 99% (dashed), and 99.9% (dash-dotted) confidence interval thresholds. The approximate 99.9% confidence intervals are: (b) $r \in (0.040, 0.054)$ [hours$^{-1}$], and (c) $\sigma^2 \in (0.084, 0.224)$ [-]. (d) Parameter-wise profile predictions for the mean (shaded) and the mathematical model simulated with the MLE (red). (e) Difference between parameter-wise profile predictions for the mean and the mathematical model simulated with the MLE. Results shown for (d,e) $r$. As there is only one parameter results for the union of the parameter-wise profile predictions are the same as (d,e).

### F Cell proliferation assay: bladder cancer cell line

In main manuscript Section 6.2 we explore an *in vitro* cell proliferation assay resulting in monolayer formation performed in [2] with NIH-3T3 fibroblast cells. Here, we explore an *in vitro* cell proliferation assay resulting in monolayer formation using the 5637 bladder cancer cell line seeded at a much larger initial density.

#### F.1 Data

The experiment is performed on tissue culture plastic with the 5637 human bladder cancer cell line (ATCC HTB-9) seeded at a density of 15,000 cells per well in a 96-well plate. The experimental duration is 136 hours (5.66 days). The plates are placed inside the IncuCyte S3 live cell imaging system (Sartorius, Goettingen, Germany) incubator (37 °C, 5% $CO_2$) [1]. Confluence measurements are obtained using automated brightfield imaging (10x objective) and processed using the IncuCyte S3 live cell imaging system (IncuCyte 2020C Rev1 Software: Basic Analyzer, phase contrast). Images are captured every two hours for the duration of the experiment. We present the confluence, $C(t) \in [0, 1]$ [-], ranging from zero to unity corresponding to 0% confluence to 100% confluent.

The 5637 human bladder cancer cell line (ATCC HTB-9) were obtained from the American Type Culture Collection (ATCC, Rockville, MD, USA) [5]. The cells were maintained in Roswell Park Memorial Institute 1640 medium (RPMI-1640; Invitrogen, Carlsbad, CA) with 10% Fetal Calf Serum (FCS, Thermo Fisher Scientific Australia, Scoresby, VIC, Australia), supplemented with 100 U/mL penicillin G (Invitrogen) in plastic tissue culture flasks in an incubator (37 °C, 5% $CO_2$). All cell lines were passaged at 2- to 3-day intervals on reaching 70% confluency using TrypLE Select (Invitrogen). Cell morphology and viability were monitored by microscopic observation and the cell line's mycoplasma-free status was confirmed using regular mycoplasma testing (Universal Mycoplasma Detection Kit, ATCC).

#### F.2 Results and Discussion

Inspecting the time evolution of $C(t)$ in this two-dimensional cell proliferation assay we observe biphasic population growth (Figure S8a). In contrast to the growth assay in the main manuscript (Figure 2), here $C(t)$ does not remain approximately constant in the first phase and instead we observe slow growth. In the second phase of growth $C(t)$ is reasonably described by logistic growth. Therefore, we explore

$$\frac{\mathrm{d}C(t)}{\mathrm{d}t} = \begin{cases} r_1 C(t), & 0 \leq t \leq T, \\ r_2 C(t)\left(1 - \dfrac{C(t)}{K}\right), & t > T. \end{cases} \tag{S.1}$$

As measurements of $C(t)$ are normalised, we treat $K$ as a known constant and set $K = 1$. We now seek estimates of five parameters, $\theta = (T, C(0), r_1, r_2, \sigma^2)$. Note that here there is one additional parameter, $r_1$, in comparison to the cell proliferation assay case study in Section 6.2.

Comparing the experimental data with the mathematical model simulated with the MLE, we observe very good agreement (Figure S8a). The profile likelihoods for the five parameters suggest that they are each identifiable to a 99.9% approximate confidence interval threshold (Figure S8b-f). Parameter-by-parameter prediction intervals reveal the influence of individual parameters on predictions (Figure S9). Fixing $T$ with and without setting $C(0)$ equal to the first experimental measurement does not accurately capture the temporal evolution of the experimental data (Figures S10a, S11a). Furthermore, approximate confidence intervals computed using the profile likelihoods of the single-phase models do not overlap with the corresponding intervals generated when analysing the data using the biphasic model (Figures S10, S11). Such differences result in accurate predictions of the temporal evolution of the experimental data when using the single-phase models (Figures S10, S11).

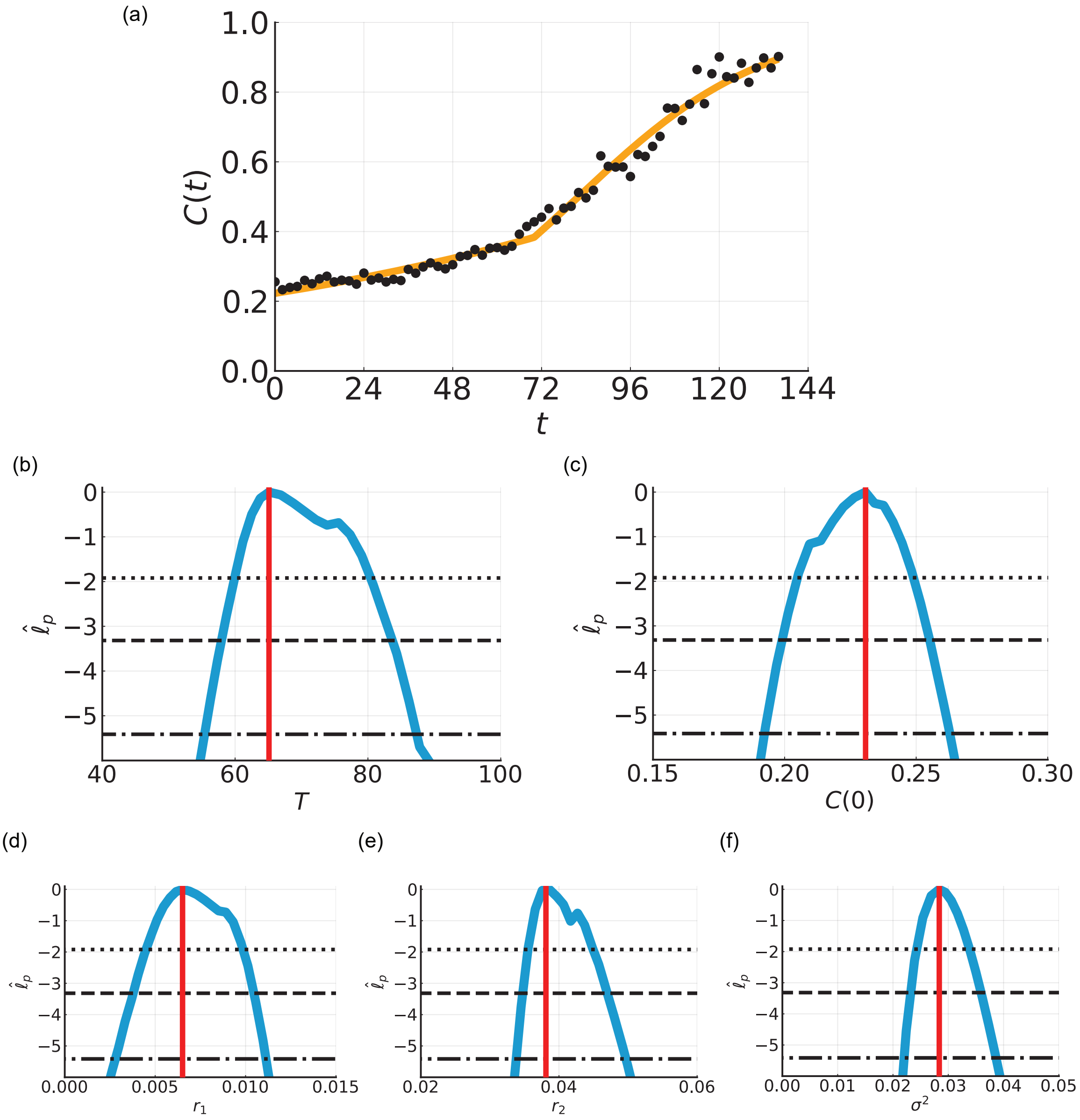


Figure S8: Biphasic population growth in a two-dimensional cell proliferation assay experiment with a bladder cancer cell line. (a) Comparison of the mathematical model simulated with the MLE (orange line) and experimental data (black circles) for the normalised cell density, $C(t)$ [-]. (b-e) Profile likelihoods for (b) $T$ [hours], (c) $C(0)$ [-], (d) $r_1$ [hours$^{-1}$], (e) $r_2$ [hours$^{-1}$], and (f) $\sigma^2$ [-] (blue) together with the MLE (vertical red line) and approximate 95% (dotted), 99% (dashed), and 99.9% (dash-dotted) confidence interval thresholds. The approximate 99.9% confidence intervals are: (b) $T \in (55.4, 87.3)$ [hours], (c) $C(0) \in (0.19, 0.26)$ [-], (d) $r_1 \in (0.003, 0.011)$ [hours$^{-1}$], (e) $r_2 \in (0.034, 0.050)$ [hours$^{-1}$], and (f) $\sigma^2 \in (0.022, 0.039)$ [-].

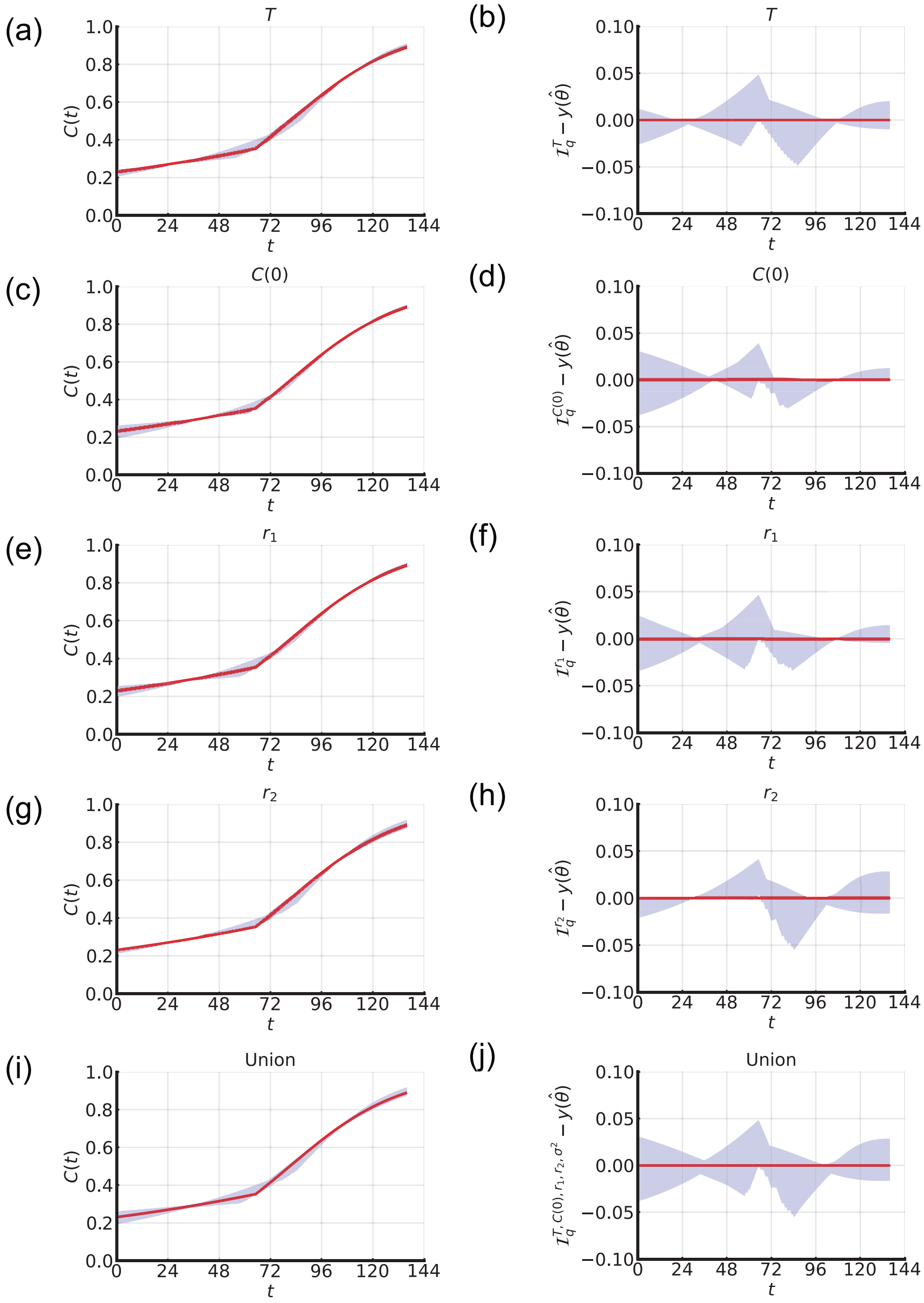


Figure S9: Parameter-wise profile predictions for two-dimensional cell proliferation assay with a bladder cancer cell line. (a,c,e,g) Parameter-wise profile predictions for the mean (shaded) and the mathematical model simulated with the MLE (red). (b,d,f,h) Difference between parameter-wise profile predictions for the mean and the mathematical model simulated with the MLE. Results shown for (a,b) $T$, (c,d) $C(0)$, (e,f) $r_1$, (g,h) $r_2$, and (i,j) the union of the parameter-wise profile predictions.

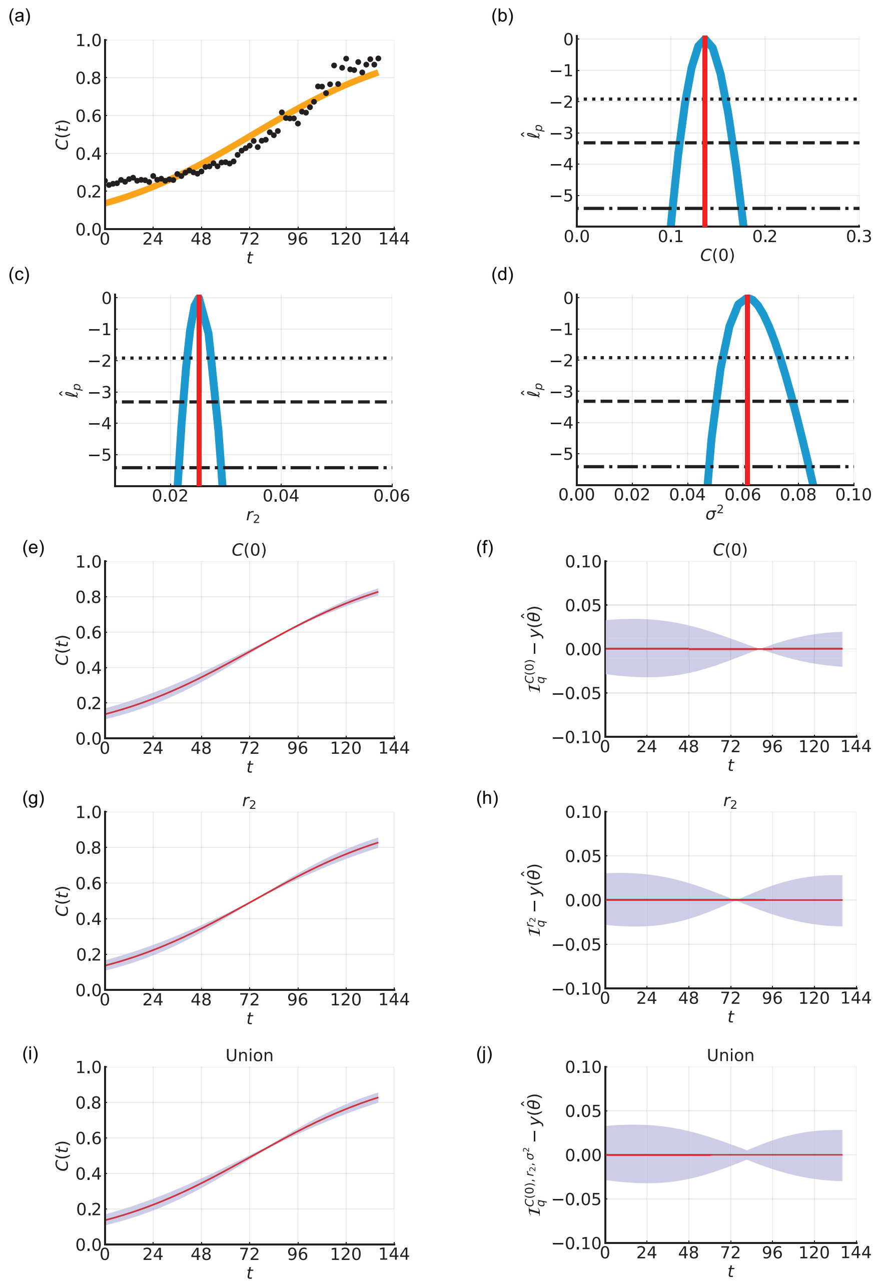


Figure S10: Biphasic population growth in a two-dimensional cell proliferation assay with a bladder cancer cell line: fixing $T = 0$. (a) Comparison of the mathematical model simulated with the MLE (orange line) and experimental data (black circles) for the normalised cell density, $C(t)$ [-]. (b-d) Profile likelihoods for (b) $C(0)$ [-], (c) $r_2$ [hours$^{-1}$], and (d) $\sigma^2$ [-] (blue) together with the MLE (vertical red line) and approximate 95% (dotted), 99% (dashed), and 99.9% (dash-dotted) confidence interval thresholds. The approximate 99.9% confidence intervals are: (b) $C(0) \in (0.10, 0.18)$ [-], (c) $r_2 \in (0.022, 0.029)$ [hours$^{-1}$], and (d) $\sigma^2 \in (0.048, 0.084)$ [-]. (e,g,i) Parameter-wise profile predictions for the mean (shaded) and the mathematical model simulated with the MLE (red). (f,h,j) Difference between parameter-wise profile predictions for the mean and the mathematical model simulated with the MLE. Results shown for (e,f) $C(0)$, (g,h) $r_2$, and (i,j) the union of the parameter-wise profile predictions.

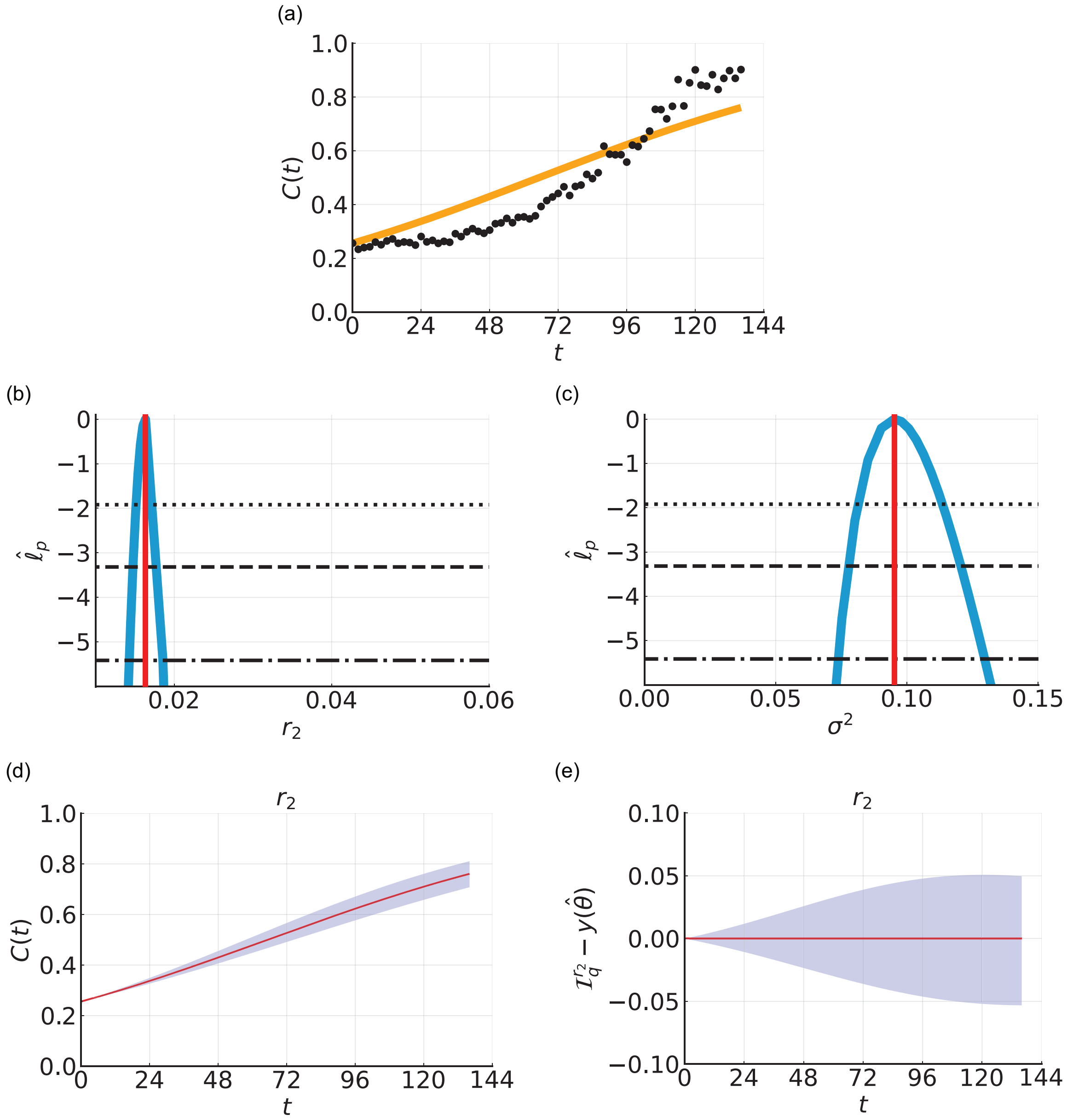


Figure S11: Biphasic population growth in a two-dimensional cell proliferation assay with a bladder cancer cell line: fixing $T = 0$ and setting $C(0)$ equal to the first experimental measurement. (a) Comparison of the mathematical model simulated with the MLE (orange line) and experimental data (black circles) for the normalised cell density, $C(t)$ [-]. (b-c) Profile likelihoods for (b) $r_2$ [hours$^{-1}$], and (c) $\sigma^2$ [-] (blue) together with the MLE (vertical red line) and approximate 95% (dotted), 99% (dashed), and 99.9% (dash-dotted) confidence interval thresholds. The approximate 99.9% confidence intervals are: (b) $r_2 \in (0.0143, 0.0186)$ [hours$^{-1}$], (c) $\sigma^2 \in (0.074, 0.130)$ [-]. (d) Parameter-wise profile predictions for the mean (shaded) and the mathematical model simulated with the MLE (red). (e) Difference between parameter-wise profile predictions for the mean and the mathematical model simulated with the MLE. (d,e) Results shown for $r_2$. As there is only one parameter results for the union of the parameter-wise profile predictions are the same as (d,e).

### G Spheroid experiment: increased experimental duration

In main manuscript Section 6.3 we analyse the three-dimensional cancer tumour spheroid experiment for $0 \leq t \leq 120$ [hours]. Here, we present results for $0 \leq t \leq 432$ [hours] (Figures S12-S13). Comparing the experimental data with the mathematical model simulated with the MLE, we observe very good agreement with small residuals (Figure S13e). By including additional data at later times we narrow approximate confidence intervals for the model parameters in the second phase, namely $r_2$ and $\mathcal{R}_2$ (Figures S13i,k). Estimates for the other parameters are very similar to results obtained when analysing $0 \leq t \leq 120$ [hours]. Parameter-wise profile predictions reveal the influence of individual model parameters on predictions (Figure S13).

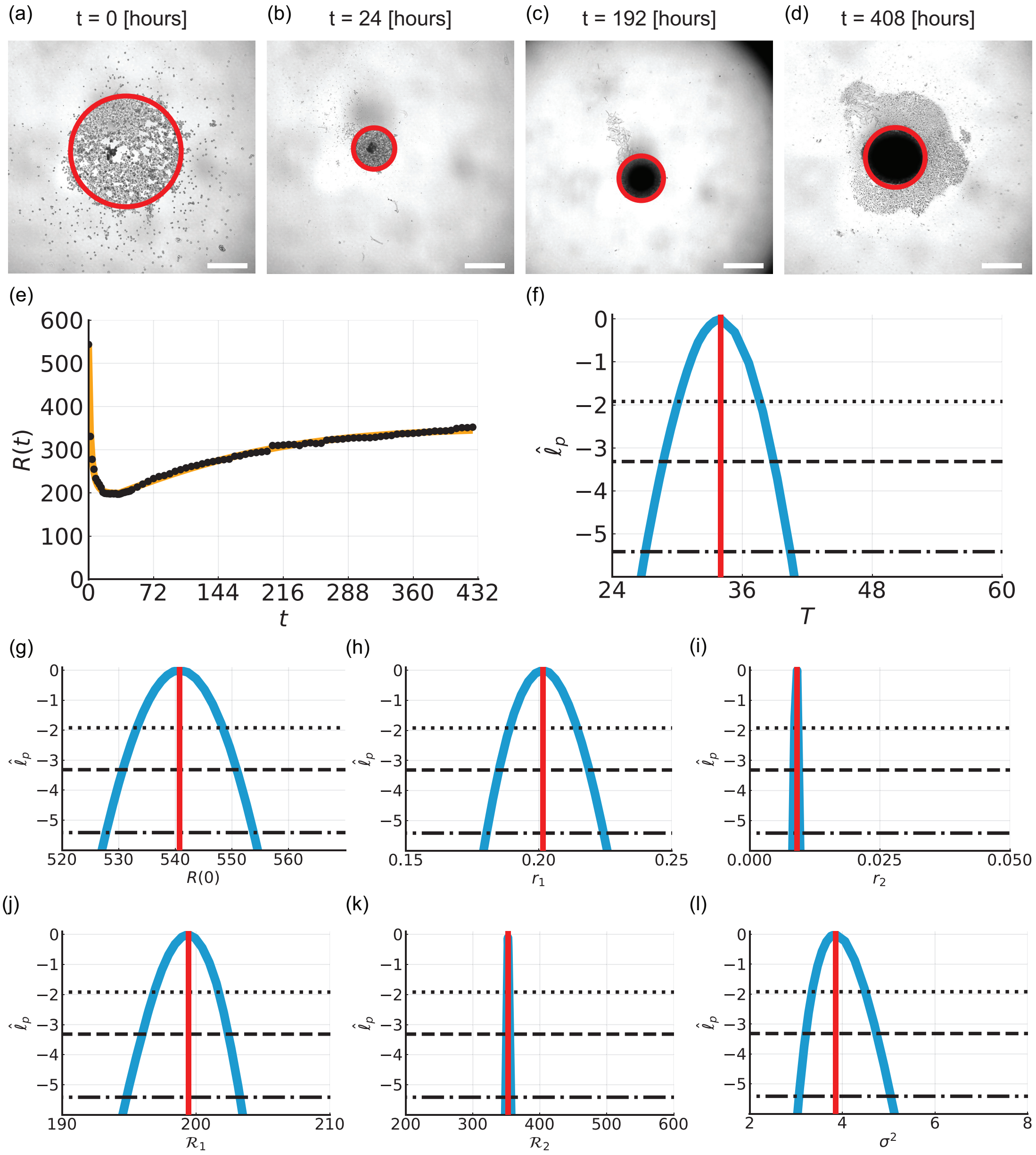


Figure S12: Biphasic population growth in three-dimensional tumour spheroid experiments for $0 \leq t \leq 432$ [hours]. (a-d) Experimental images of spheroid experiments at (a) $t = 0$ [hours], (b) $t = 24$ [hours], (c) $t = 192$ [hours], and (d) $t = 408$ [hours]. Scale bar is 400 µm and red circle shows approximate equivalent area. At early times cells migrate and adhere to form spheroid. At later times the spheroid grows as a compact mass. (e) Comparison of the mathematical model simulated with the MLE (orange line) and experimental data (black circles) for the equivalent radius, $R(t)$ [µm]. (f-l) Profile likelihoods for (f) $T$ [hours], (g) $R(0)$ [µm], (h) $r_1$ [hours$^{-1}$], (i) $r_2$ [hours$^{-1}$], (j) $\mathcal{R}_1$ [µm], (k) $\mathcal{R}_2$ [µm], and (l) $\sigma^2$ [-] (blue) together with the MLE (vertical red line) and approximate 95% (dotted), 99% (dashed), and 99.9% (dash-dotted) confidence interval thresholds. The approximate 99.9% confidence intervals are: (f) $T \in (27.1, 40.5)$ [hours], (g) $R(0) \in (527.8, 553.8)$ [µm], (h) $r_1 \in (0.180, 0.224)$ [hours$^{-1}$], (i) $r_2 \in (0.008, 0.009)$ [hours$^{-1}$], (j) $\mathcal{R}_1 \in (194.8, 203.3)$ [µm], (k) $\mathcal{R}_2 \in (348.8, 356.8)$ [µm], and (l) $\sigma^2 \in (3.06, 5.04)$ [-].

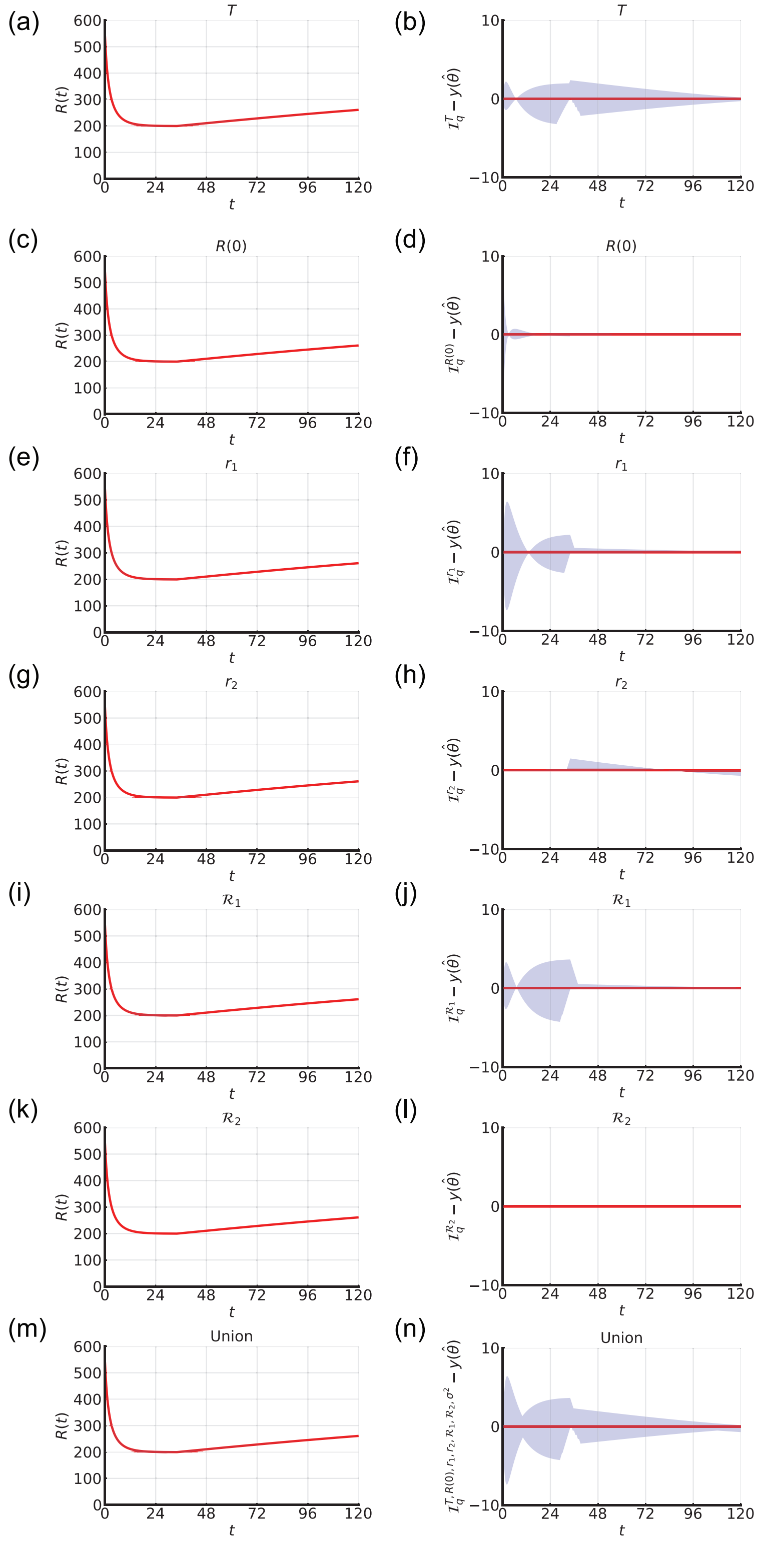


Figure S13: Parameter-wise profile predictions for three-dimensional cancer tumour spheroid experiment ($0 < t < 432$ [hours]). (a,c,e,g,i,k,m) Parameter-wise profile predictions for the mean (shaded) and the mathematical model simulated with the MLE (red). (b,d,f,h,j,l,n) Difference between parameter-wise profile predictions for the mean and the mathematical model simulated with the MLE. Results shown for (a,b) $T$, (c,d) $R(0)$, (e,f) $r_1$, (g,h) $r_2$, (i,j) $\mathcal{R}_1$, (k,l) $\mathcal{R}_2$, and (m,n) the union of the parameter-wise profile predictions.
